# Persistent climate maladaptation decreases performance of a keystone tree species (*Quercus lobata*)

**DOI:** 10.64898/2026.02.06.704494

**Authors:** Alexander R. B. Goetz, Marissa E. Ochoa, Berenice Badillo, Jessica W. Wright, Victoria L. Sork

## Abstract

Maladaptation in foundational tree species can undermine ecosystem stability, depending on its current magnitude and future extent under climate change. Using a range-wide common garden study of valley oak (*Quercus lobata*), we tested whether historical warming has already produced climate–fitness mismatches, whether climatic extremes intensify these mismatches, and how these patterns can inform management. Half-sib families of 12-year-old trees had the highest growth and survival under summer temperatures cooler than their home climates, demonstrating persistent maladaptation to contemporary temperature maxima. This mismatch intensified during hotter years and imposed lasting performance effects. Simulated assisted gene flow using seeds of high-performing maternal families from sites warmer than the planting sites yielded greater projected performance than climate matching alone. These findings provide phenotypic evidence that maladaptation is constraining a keystone tree species and suggest that genotypes pre-adapted to future conditions can be identified throughout the species range to improve resilience under further warming.

## Introduction

Rapid climate change is increasing the risk that organisms will become maladapted to their local environments (Jump & Peñuelas 2005; Derry *et al*. 2019; Kooyers *et al*. 2025), threatening ecosystem function worldwide (Intergovernmental Panel on Climate Change 2023). Maladaptation—the distance between optimal and actual fitness (e.g., Brady *et al*. 2019b)—can occur if climate change outpaces selection and dispersal (Carter 1996; Davis *et al*. 2005; Aitken *et al*. 2008; Thuiller *et al*. 2008; Gougherty *et al*. 2021; Rellstab 2021). Many studies predict increasing maladaptation under climate change (e.g., Anderson & Wadgymar 2020; Rellstab 2021; Xu & Prescott 2024), but the magnitude of maladaptation and its consequences for population performance remain poorly quantified. This gap limits our ability to determine whether maladaptation poses a substantial threat to species persistence and whether affected populations would benefit from management intervention. Reduced fitness in ecologically important species may lead to cascading and irreversible ecosystem consequences (Valiente-Banuet *et al*. 2015; Ellison 2019), yet some degree of maladaptation is expected even in stable environments and may have limited demographic effects (Brady *et al*. 2019a; Derry *et al*. 2019; Leites & Benito Garzón 2023). Determining not only whether maladaptation occurs, but how much it affects population performance, is therefore essential for evaluating strategies to mitigate its effects.

A useful way to assess whether species can keep pace with rapid human-driven climate change is to first consider how they responded to slower climatic shifts in the past. For some species, natural selection and gene flow may not keep pace with rapid future climate change (e.g., St. Clair & Howe 2007; Frank *et al*. 2017b; Anderson *et al*. 2025), producing an adaptational lag that shifts populations from their optimal climates (Davis *et al*. 2005; Hendry & Gonzalez 2008; Borgman *et al*. 2015). Trees may be especially susceptible to this lag because their long generation times slow evolutionary response (Frank *et al*. 2017a; Browne *et al*. 2019; Gougherty *et al*. 2021; Rellstab 2021; Benomar *et al*. 2022; Schmeddes *et al*. 2024) and their limited seed dispersal capacity constrains range shifts (Davis & Shaw 2001; Aitken *et al*. 2008; Corlett & Westcott 2013; Merlin *et al*. 2018). Many tree species are projected to become maladapted to their new environments as climate warming accelerates (e.g., Frank *et al*. 2017a; Chakraborty *et al*. 2019; Diamond & Martin 2020; Rellstab 2021), while others are already poorly adapted to contemporary climates (Rehfeldt *et al*. 2001; Savolainen *et al*. 2004; St. Clair & Howe 2007; Browne *et al*. 2019; Foest *et al*. 2025). Because many species struggled to keep pace even with gradual climatic shifts in the past, it is unlikely they will track the far more rapid changes ahead, raising concerns for forest resilience and long-term ecosystem stability. Quantifying the degree to which species have adapted, or failed to adapt, to current environments can therefore inform predictions of how they may perform under dynamic future conditions.

Under fluctuating climates, the degree of maladaptation may vary through time (Brady *et al*. 2019a; Wolkovich & Donahue 2021). Interannual variability in climate can alter selective pressures on trees (Anderson 2016; Girardin *et al*. 2016; Girardin *et al*. 2021), and climate change is increasing the frequency and severity of weather extremes (Wang *et al*. 2017). Such extremes may amplify maladaptation if trees cannot tolerate or recover from them (Lloret *et al*. 2011; Benito Garzón *et al*. 2013a; Depardieu *et al*. 2020). Yet inferences about climatic adaptation in trees are often constrained by the short duration of experiments, which may fail to capture a realistic range of weather conditions (Bowman *et al*. 2013; Aspalter *et al*. 2025). Tracking fitness-climate relationships across years can reveal whether short-term weather extremes influence the magnitude of maladaptation. In addition, relationships between climate and fitness are typically tested using seedlings or young trees (e.g., Gellie *et al*. 2016; Browne *et al*. 2019; Butnor *et al*. 2019; Etterson *et al*. 2020), which may misrepresent patterns in later life stages (Germino *et al*. 2019; Ramírez-Valiente *et al*. 2022). Responses of young trees to the environment inferred from experiments may be shaped by maternal effects (Roach & Wulff 1987) or planting shock (Close *et al*. 2005), temporarily masking true climate–fitness relationships. Alternatively, acclimation to garden conditions may reduce apparent maladaptation as trees age (e.g., Martínez-Sancho *et al*. 2025). Consequently, long-term studies that track tree performance across years and life stages are essential to distinguish transient responses from persistent patterns and to accurately characterize the degree of maladaptation in tree populations.

Predicting future maladaptation requires robust measures of how the environment affects current fitness. A powerful and increasingly widespread method to forecast the effects of climate maladaptation on populations across time and space is genomic offset modeling (Capblancq *et al*. 2020), which is done by associating genomic data with current and future climatic gradients. However, this method assumes local adaptation to current climate (Lind *et al*. 2024; Lotterhos 2024a) and is therefore limited to predicting relative changes in adaptedness, or the degree to which organisms are fit to the environment in which they grow, *sensu* Allard (1988). Genomic predictions therefore have a major limitation: Even if their predictions are validated empirically (e.g., Archambeau *et al*. 2024; Verrico *et al*. 2026), they cannot tell us how detrimental it is for a species to be maladapted in the future (Lotterhos 2024a; Fitzpatrick *et al*. 2026). By contrast, models based on phenotypic data from common garden studies can predict actual fitness consequences (e.g., Carter 1996; Etterson *et al*. 2020), which means we can determine the degree to which populations are currently adapted to their environments and thus predict how future changes will affect actual fitness components (Carter 1996; Wang *et al*. 2010; Benito Garzón *et al*. 2019; Huxman *et al*. 2022; Leites & Benito Garzón 2023). Thus, we can robustly test the degree to which maladaptation will affect future populations and explore whether hypothetical assisted gene flow may improve resilience.

The overarching goal of this study is to test the degree to which maladaptation to climate affects a widespread California keystone tree species, valley oak (*Quercus lobata* Née). Valley oak populations are threatened by habitat reduction (Kelly *et al*. 2005), recruitment limitations (Tyler *et al*. 2006; Zavaleta *et al*. 2007), and predicted range shifts due to climate change (Kueppers *et al*. 2005; Zavaleta *et al*. 2007; Sork *et al*. 2010; McLaughlin & Zavaleta 2012), all of which would be intensified by sustained climate maladaptation. We used a 12-year longitudinal dataset generated by a long-term provenance study to determine the extent of maladaptation in juvenile trees, and to test whether its strength varies with fluctuating temperature. Previous work in the same common gardens when the trees were five years old (Browne *et al*. 2019) allows us to compare evidence of maladaptation across two time points and test whether maladaptation increases or decreases as trees age. We first analyzed growth and survival of 3,415 half-sib offspring of 628 maternal trees planted in two common gardens, which allowed us to quantify climate–fitness relationships of families originating from a large portion of the species’ range. Then, we used these climate–fitness relationships to predict future changes in performance. We addressed three questions: (1) How have past changes in climate resulted in maladaptation? (2) How much do short-term weather extremes influence maladaptation? (3) Can simulations of assisted gene flow demonstrate its potential to mitigate the effects of maladaptation? By integrating long-term performance data with climate variation, this study provides evidence that maladaptation in valley oak affects individual growth and survival and that this knowledge can inform future management strategies aiming to reduce the impact of maladaptation.

## Materials and methods

### Study species

Valley oak (*Quercus lobata* Née) is a long-lived (up to 500 or more years), winter-deciduous tree species endemic to woodlands, savannas, and riparian zones of the California Floristic Province, 0—1700m above sea level (Pavlik *et al*. 1991). Though its historical range prior to European colonization likely covered much more of the state, up to 95% of that range has already been lost to land conversion (Kelly *et al*. 2005). Furthermore, evidence shows that current valley oak populations are adapted to temperatures from the Last Glacial Maximum rather than the recent past, but the degree of apparent maladaptation differs across genetic lines (Browne *et al*. 2019). Finally, shifts in local climatic niches of valley oak populations are predicted to outpace gene flow, limiting its ability to adapt (Sork *et al*. 2010).

### Experimental design

This study utilizes two long-term common gardens established by JW Wright and VL Sork in 2012 as part of a long-term provenance test of valley oak (described in Delfino Mix *et al*. 2015). To establish the provenance test, 11,000 open-pollinated acorns were collected from 674 adult trees at 95 localities (AKA provenances) across the species range of valley oak (Figure 1, black dots). Acorns were collected from 5-8 trees per locality. Twelve acorns per tree, comprising half-sib families, were then germinated in a greenhouse.

**Figure 1:**
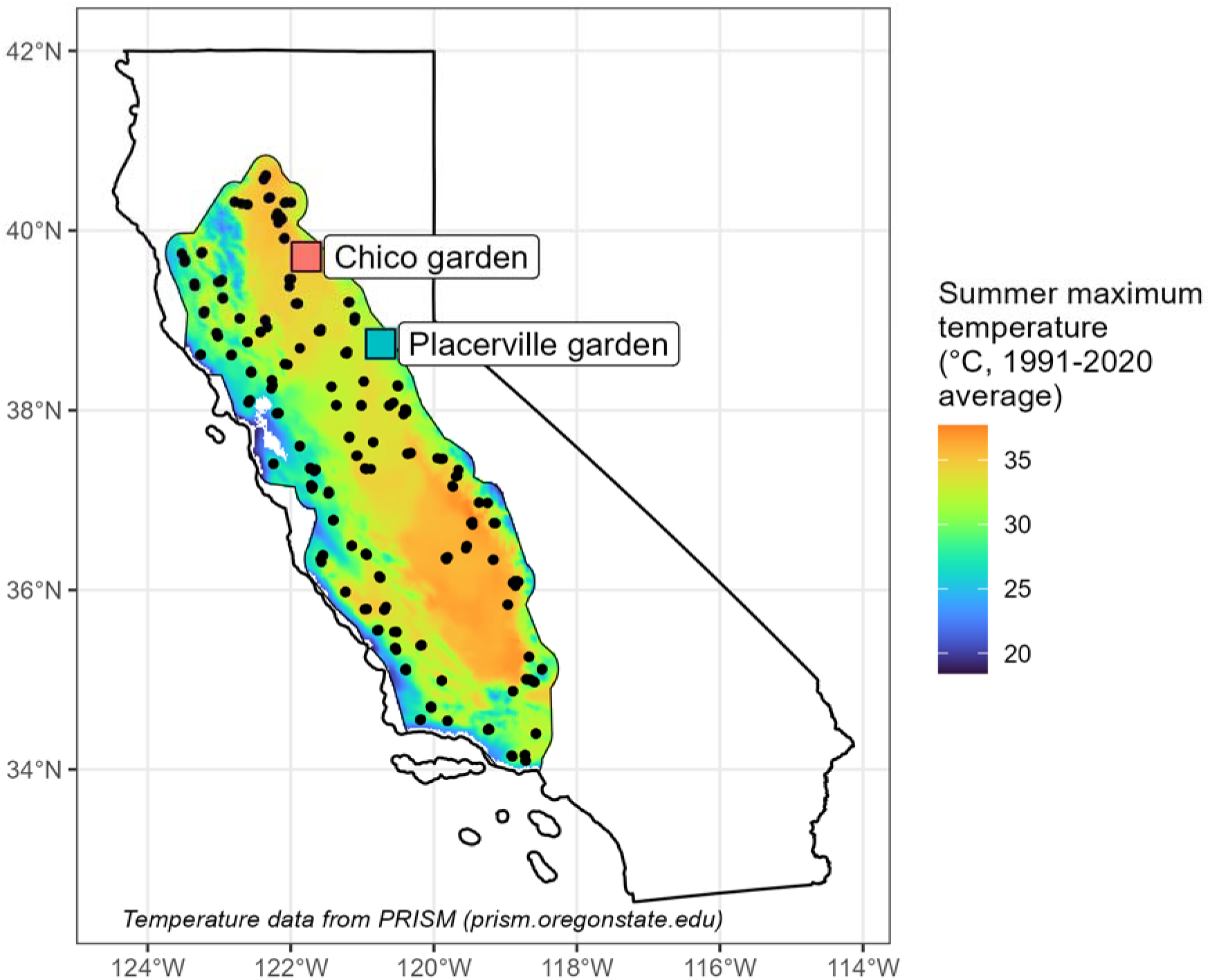
Mid-point localities (black circles, n = 95) of sets of 3-5 maternal *Q. lobata* trees used as seed sources for 628 families planted into two common gardens (labeled squares). Outline represents the theorized current species range based on the distribution of living trees (Davis *et al*. 1998). Labeled squares represent the two common garden locations (red: Chico Seed Orchard, Chico, CA; blue: Institute of Forest Genetics, Placerville, CA). Background color represents 1991-2020 average of maximum yearly summer temperature (data from PRISM: prism.oregonstate.edu). The distribution of summer and winter temperatures across localities is provided in Figure S2.

This study treats localities and trees as genetically independent of one another because population structure in valley oak is very low (Sork *et al*. 2010; Gugger *et al*. 2013; Ashley *et al*. 2015) and individuals growing at least 100m apart are unlikely to be closely related (Dutech *et al*. 2005). Therefore, while we use locality as a blocking factor to account for spatial autocorrelation in environmental characteristics, we do not treat localities as distinct genetic subpopulations. Instead, we measure clinal effects of climate on quantitative traits at the family level (Callaham 1963; Mátyás 1996).

Following one year of growth in a greenhouse and one year of growth in lath houses, 3-5 seedlings from each family were planted into each of two common gardens (“Placerville garden”: Placerville, CA, 822m above sea level; “Chico garden”: Chico, CA, 70m above sea level) in 2015. Both gardens have intermediate climates relative to the species distribution (Delfino Mix *et al*. 2015), but the Chico garden is hotter and drier than the Placerville garden. To allow for testing of genotype-by-environment (G x E) effects on phenotypes, seedlings from all families were planted in each garden. To ensure tree survival for measurement of phenotypes, the Placerville garden was irrigated from 2015—2019 and the Chico garden was irrigated in all years. In 2021, according to the original study plan to reduce inter-tree competition as trees grew, roughly half of the individuals at each garden were thinned. As of 2024, the two gardens contained a total of 3,415 surviving juvenile trees from 628 families (Placerville: 1,673 trees from 623 families; Chico: 1,742 trees from 626 families), with 1-5 surviving individuals per family in each garden. Due to multiple environmental differences between gardens (e.g., elevation, hydrology, seasonality, and duration of irrigation), our analyses treat them categorically as two distinct environments instead of “warm” and “cool” treatments. We measured height and survival of all trees from 2014-2024 (refer to Appendix S1 and Table S1 for details).

### Testing for effects of temperature transfer distance on fitness components

To answer Q1, we tested the effects of climate on components of fitness in valley oak. As background to our analyses, we first tested whether a standardized metric of tree height, height′ (Appendix S2) was genetically differentiated among families and years (Appendix S3). To compare the climates of the maternal seed sources to those of the two gardens, we calculated transfer distances (Eriksson et al. 1980, Prescher 1986) based on maximum summer temperatures and mean winter temperatures for each family in each year, taken as the difference between contemporary (2014-2024) temperatures in each garden and 30-year averages of recent-past (1951-1980) temperatures at the maternal origins (Appendix S4). We used 1951-1980 averages for the origin climates, taken from the Basin Characterization Model (Flint *et al*. 2013) at 270m resolution, because they are more similar to the climates in which the maternal trees (which may be up to 300 years old or more) would have established than modern temperatures, especially because climate was more stable across California prior to the 1990s (LaDochy & Witiw 2023). Maximum summer temperature in 1951-1980 at the study sites was correlated with estimated maximum summer temperature for the 1700-1800 and 1900-2000 centuries (Figure S1; Karger et al. 2023); we use climatic data from the BCM in our analyses due to its high spatial resolution. We examined maximum summer temperatures because they (among other climate variables) have been shown to constrain valley oak distribution (Kueppers et al. 2005, Sork et al. 2010, McLaughlin and Zavaleta 2012, Gugger et al. 2013). We examined mean winter temperatures because they reflect selective pressure imposed by freezing damage (Ramírez-Valiente *et al*. 2022). Summer and winter temperatures are also both predicted to increase in the future (Wang et al. 2017). In addition, because summer maximum temperature and mean winter temperature showed a small but significant quadratic relationship in our study sites (Figure S2), we tested the effects of mean winter temperature on fitness as well.

We tested the effects of temperature transfer distances on components of fitness, which we measured as two separate variables: 1) Cumulative and annual relative growth rates (RGR) of individuals from 2014-2024 (Appendix S5), and 2) relative fitness, calculated as average height′ in 2024 × percent survival of each family from 2014-2024 (Appendix S6). First, we used linear mixed models to estimate variance components, from which we calculated quantitative trait differentiation (Q_ST_) values and narrow-sense heritability (Falconer & Mackay 1983) for cumulative relative growth rate. We then further analyzed the data by fitting generalized additive models (GAM; Hastie & Tibshirani 1986; Wood *et al*. 2015). We used GAMs so that we could test the fitness-climate relationship without assuming its shape *a priori*. We interpreted roughly quadratic relationships as evidence of local adaptation (Callaham 1963) whereas we interpreted roughly linear relationships as evidence of a “shifted” distribution suggesting adaptational lag (Browne *et al*. 2019). Models were fitted using the following formulas (refer to Appendix S7 for details):

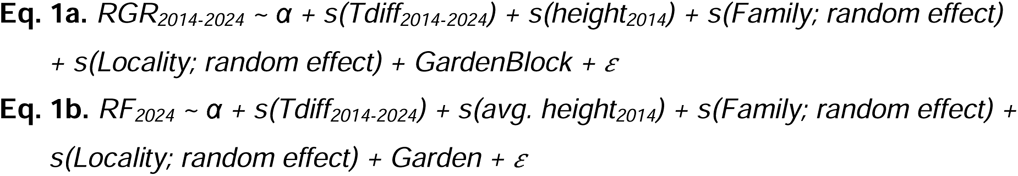

where α is the intercept and is the error term; RGR is relative growth rate; RF is relative fitness; Tdiff_2014-2024_ is temperature transfer distance between the source site average (1951-1980 mean) and each garden (2014-2024 mean). Terms within an s() parenthetical are smooth terms modeled using cubic regression splines.

### Testing for changes in intensity of maladaptation among years

To test whether the degree of maladaptation varied across years (Q2), we used a longitudinal GAM approach to analyze the extent to which growth rates of trees from different origin climates vary across years. We fit the GAMs using the same software methods as done for the cumulative models (Appendix S7). The independent variable of interest was the interaction between study year and temperature difference (modeled as a hierarchical cubic regression spline with a group-specific smoother for the study year effect; Pedersen *et al*. 2019). We also included both 2014 tree height and tree height in the previous study year to account for the fact that growth rate inherently decreases with increasing biomass (Rees *et al*. 2010). The model was fitted using the following formula:

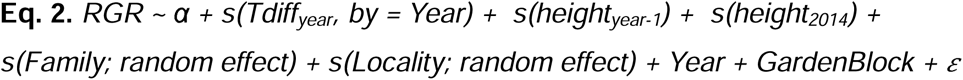

where α is the intercept and is the error term; Tdiff_year_ is temperature transfer distance within each year. Terms within an s() parenthetical are smooth terms modeled using cubic regression splines.

To determine whether hotter annual temperatures reduced relative growth rates irrespective of trees’ source climates, we tested the effect of annual maximum temperature on average RGR across all trees in both gardens using a linear regression. To test whether trees from different origins differed in their resilience to hotter temperatures (Lloret *et al*. 2011), we examined whether hotter study years were associated with lower RGR among families from distinct climate origins in each garden. To determine if the previous year’s temperature influenced growth in the next year, we included the interaction between the origin temperature and the previous year’s temperature (ANCOVA; functions lm() and anova() in R package stats). We excluded the first two study years (2015 and 2016) because all families exhibited very high RGR in these years, likely due to outplanting or ontogenetic effects. As in the other RGR analyses, we also included the effect of planting block to account for known environmental differences within gardens. Finally, we tested the effects of phenology on growth to determine whether differences were driven by growing season length (Appendix S8).

### Extrapolating theoretical fitness maxima

To aid in interpreting the climate-fitness relationship (Q1, Q2), we extrapolated the temperature transfer functions beyond the minimum observed transfer distances using Gaussian process regression. Gaussian process regression extrapolates model fits under the assumption that the modeled variable follows a normal distribution (Rasmussen & Williams 2006), which models the relationship between environment and fitness under local adaptation (Savolainen *et al*. 2007). Thus, if a climate-fitness relationship does not show a peak (e.g., Browne *et al*. 2019), extrapolating beyond the observed limit allows us to estimate where the peak would occur if more extreme transfer distances could be tested. This extrapolation is not an empirical test, but it allows us to estimate the underlying climate-fitness relationship that would lead to our observed data.

To estimate the transfer distance that would result in a fitness peak, we extrapolated from the GAM-fitted values of relative growth rate and relative fitness. We fitted Gaussian process equations to each model, estimating mean and variance based on the shape of the curve, and predicted values out to temperatures where the origin site would have a maximum summer temperature 12°C warmer than the common gardens, roughly 8°C beyond the greatest negative transfer distance observed in the study. We used 12°C as the prediction limit because it was approximately equal to the temperature transfer distance between the present-day Chico garden and the average temperature of all maternal origin sites during the Last Glacial Maximum, so it represented a realistic range of the shift in temperature between the Last Glacial Maximum and the present. Gaussian process regression was fitted using function gpkm() with a Matern 5/2 kernel in R package ‘GauPro’, version 0.2.13 (Erickson 2024).

### Simulating the effects of assisted gene flow on future fitness

To test how maladaptation to changing climates will affect future populations of valley oak and whether hypothetical assisted gene flow efforts can bolster future forest resilience (Q3), we modeled three future scenarios of range-wide relative fitness distribution. The scenarios were: 1) local trees remaining in place (“local acorns”), 2) selecting seed sources whose current climates best match projected future conditions (“climate matching”), and 3) first identifying seed sources through climate matching and then selecting the highest performers (“performance-based climate matching”). Performance-based climate matching therefore combines climate matching (Aitken *et al*. 2008) with a classical tree improvement approach (White 1987) to maximize performance under warmer future conditions.

To predict future relative fitness, we first constructed a series of rasters representing baseline distributions of relative fitness under the historical climates to which populations are adapted (phenotype-environment association; Appendix S9). These models allowed us to set up each scenario based on hypothetical present-day management action informed by adaptation to historical conditions and then observe its effects after 50-70 years of rapid climate change under a Relative Concentration Pathway 4.5 stabilization scenario (Intergovernmental Panel on Climate Change 2023).

To model Scenario 1 (local genotypes remaining in place), we used the null fitness surface calculated from Eq. 1b. To model Scenario 2 (seeking preadapted seed sources solely based on site temperature as a criterion), we randomly selected a common garden tree from a warmer site than the focal pixel within a 25-km radius. To model Scenario 3 (seeking preadapted seed sources based on site temperature and performance), we selected the tree with the highest performance value within a 25 km radius, again from a site warmer than the focal pixel. To avoid circularity caused by selecting top performers based on their 2024 relative fitness, which was used to fit the transfer functions, we instead selected top performers based on their 2017 relative fitness (when seedlings were five years old). Thus, Scenario 3 simulated the use of seedling data to make fitness predictions. We chose 25km as a conservative radius for hypothetical seed transfer due to concerns about unintended effects of long-distance transfer (McKay *et al*. 2005; O’Neill *et al*. 2014).

To predict the consequences of each scenario across the species range, we calculated a T_diff_ raster representing in-place temperature change between past and future (2071-2100 average of maximum summer temperature – 1961-1990 average of maximum summer temperature). Projected future climate data were taken from the AdaptWest 8-model ensemble projections (AdaptWest Project 2022; Mahony *et al*. 2022). See Appendix S9 for further details.

Finally, we predicted future relative fitness for each pixel using the relationship between T_diff_, baseline relative fitness, and future relative fitness, fitted with Eq. 1b. To generalize the model across the species range, we averaged the fitness data across source localities and gardens and removed those terms from the model. We then projected potential fitness distributions of hypothetical 12-year-old valley oak trees in year 2100 under an RCP 4.5 climate change pathway. Finally, we calculated the percent change in fitness relative to baseline conditions across the species range.

## Results

### Background

Height of trees, a component of fitness, showed significant genetic differentiation among families (ANOVA; df = 630, 2417; P < 0.001), and gardens (ANOVA, df = 1, 2417; P < 0.001). The family × garden interaction term was not significant (ANOVA, df = 620, 2417; P = 0.24) indicating that the same genotypes tended to grow either better or worse in both gardens (Table S2). Cumulative relative growth rate was highly genetically differentiated in both gardens, but heritability was much higher at the Placerville garden than at the Chico garden and additive genetic variance was low at both gardens (Table 1). In addition, relative growth rate significantly differed among families across years, providing evidence of interannual variation in growth (Table S3).

**Table 1:** Estimated variance components (V_A_: additive genetic variance, V_P_: total phenotypic variance), quantitative trait differentiation (Q_ST_), and narrow-sense heritability (*h^2^*) of cumulative relative growth rates, 2014-2024, in two common gardens.

| **Garden** | **V<sub>A</sub>** | **V<sub>P</sub>** | **Q<sub>ST</sub>** | ***h<sup>2</sup>*** |
|---|---|---|---|---|
| **Placerville** | 0.0014 | 0.0039 | 0.59 | 0.35 |
| **Chico** | 0.0002 | 0.0023 | 0.88 | 0.07 |

### Evidence of maladaptation to climate in 12-year-old trees

Based on relative growth rates and relative fitness, families of 12-year-old valley oak trees showed significant maladaptation to summer temperatures (Figure 2A, 2C). When examining summer maximum temperature differences, we observed faster growth rates of individuals (Figure 2A) and higher relative fitness per family (Figure 2C) in trees originating from warmer source sites than the common gardens. The theoretical trait optima of growth rate and relative fitness both occurred closer to -5.2°C, the summer transfer distance from the average temperature during the Last Glacial Maximum (approximately 21,000 years ago; (Karger *et al*. 2017; Karger 2025), than to a transfer distance of zero (Figure 2A, 2C), which is consistent with the hypothesis of adaptational lag. Likewise, relative growth rate and relative fitness showed further declines under a predicted future summer temperature increase of 1.8°C (Figure 2A, 2C). Model summaries and cross-validation RMSEs are provided in Tables S4-S7.

**Figure 2:**
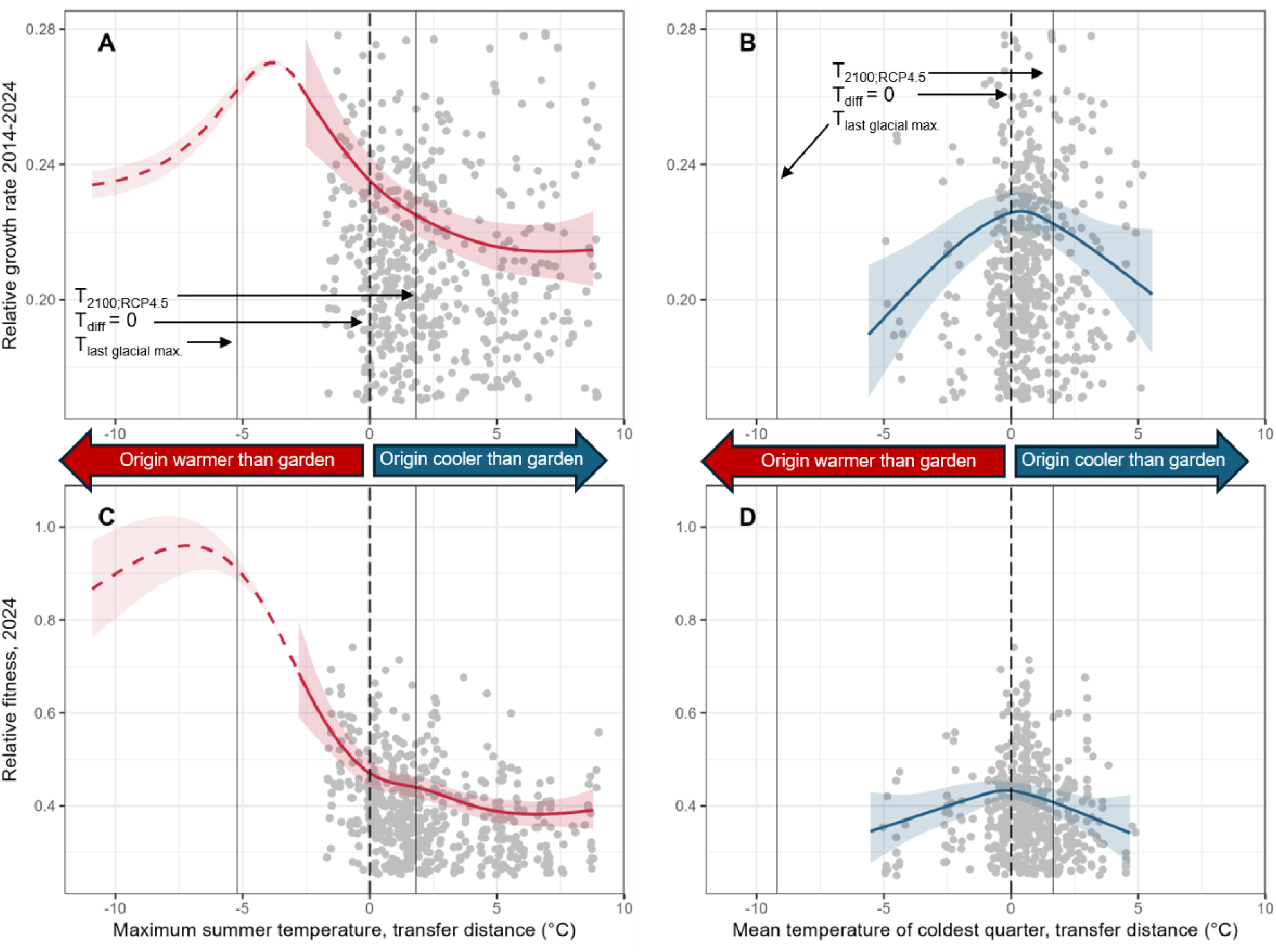
Evidence of maladaptation to summer maximum temperatures and local adaptation to winter mean temperatures in 12-year-old *Q. lobata* trees growing in two common gardens. A-B. Cumulative relative growth rate (2014-2024). **C-D.** Relative fitness (growth × survival) as of 2024. Solid curves denote generalized additive models fit to empirical data, and dashed curves denote extrapolation of the models using Gaussian process regression. Shaded regions represent 95% confidence intervals. Red curves (**A, C**) are models of summer maximum temperature and blue curves (**B, D**) are models of winter mean temperature. The peak in each extrapolated curve represents the theoretical species-wide trait optimum. Grey points are family average RGRs across both sites (**A, B**) or family relative fitness values (**C, D**). Vertical dashed line in each plot denotes where source temperature equals garden temperature; vertical solid lines denote estimated average temperature during the Last Glacial Maximum (21 kya; Wang et al. 2012) and predicted average temperature in year 2100 under a stabilization climate scenario (RCP 4.5; AdaptWest Project 2022). The area to the left of the zero line contains individuals from warmer historical climates than the garden, while the area to the right contains individuals from cooler historical climates. See Tables S4-S7 for statistical summaries.

By contrast, we observed evidence of local adaptation to winter temperatures (Figure 2B, 2D). Trees from origins with similar winter temperatures to the common gardens had the highest relative growth rates and relative fitness values, with decreasing values of both growth metrics in trees further from T_diff_ = 0 (Figure 2B, 2D). Growth would therefore be projected to decrease slightly under the predicted warmer future winter temperature increase of 1.3°C (Intergovernmental Panel on Climate Change (IPCC) 2023; Figure 2A, 2C, solid vertical lines). Model summaries and cross-validation RMSEs are provided in Tables S8-S9.

### Variation in maladaptation among years

Because the relationship between temperature and growth is (a) variable across years and (b) does not relax over time, our results provide no evidence of either short-term plasticity or long-term acclimation to the garden environment, either of which could mitigate maladaptive responses. Instead, the effect of temperature on growth varied among years (Figure 3A, Table S10), with a more extreme relationship between temperature and growth in hotter years (Figure 3A, Table S10). Higher annual maximum temperatures were significantly associated with lower growth, with a consistently linear trend showing that any increase beyond the lowest observed Tmax (∼33°C) reduced annual growth (Figure 3B, Table S10). Furthermore, garden temperature in the year prior to the growing season also significantly predicted variation in relative growth rates (Figure 3C, Table S11), indicating that the negative effect of a hot year persists in future years. While the slope of the transfer distance–growth relationship varied over time, it remained consistently negative from years 5-12 (2017-2024; Figure 3A). This pattern means that individuals from warmer sites than the common gardens consistently had the highest growth rates, even in cooler years, leading to a widening gap in relative fitness (Figure 3C). Trees were larger at the Chico garden than the Placerville garden due to exceptional early growth in the first two study years (2015 and 2016); from 2017 onward, relative growth rates have been similar at both gardens (Figure 3C). Therefore, we find that short-term hotter temperatures further reduce the adaptedness of all genotypes, but especially genotypes originating from cool climates to the garden environment. See Tables S10 and S12 for model summaries and cross-validation RMSEs. Finally, we did not find that phenology was a primary driver of differences in growth among families or years (Tables S13-S14).

**Figure 3:**
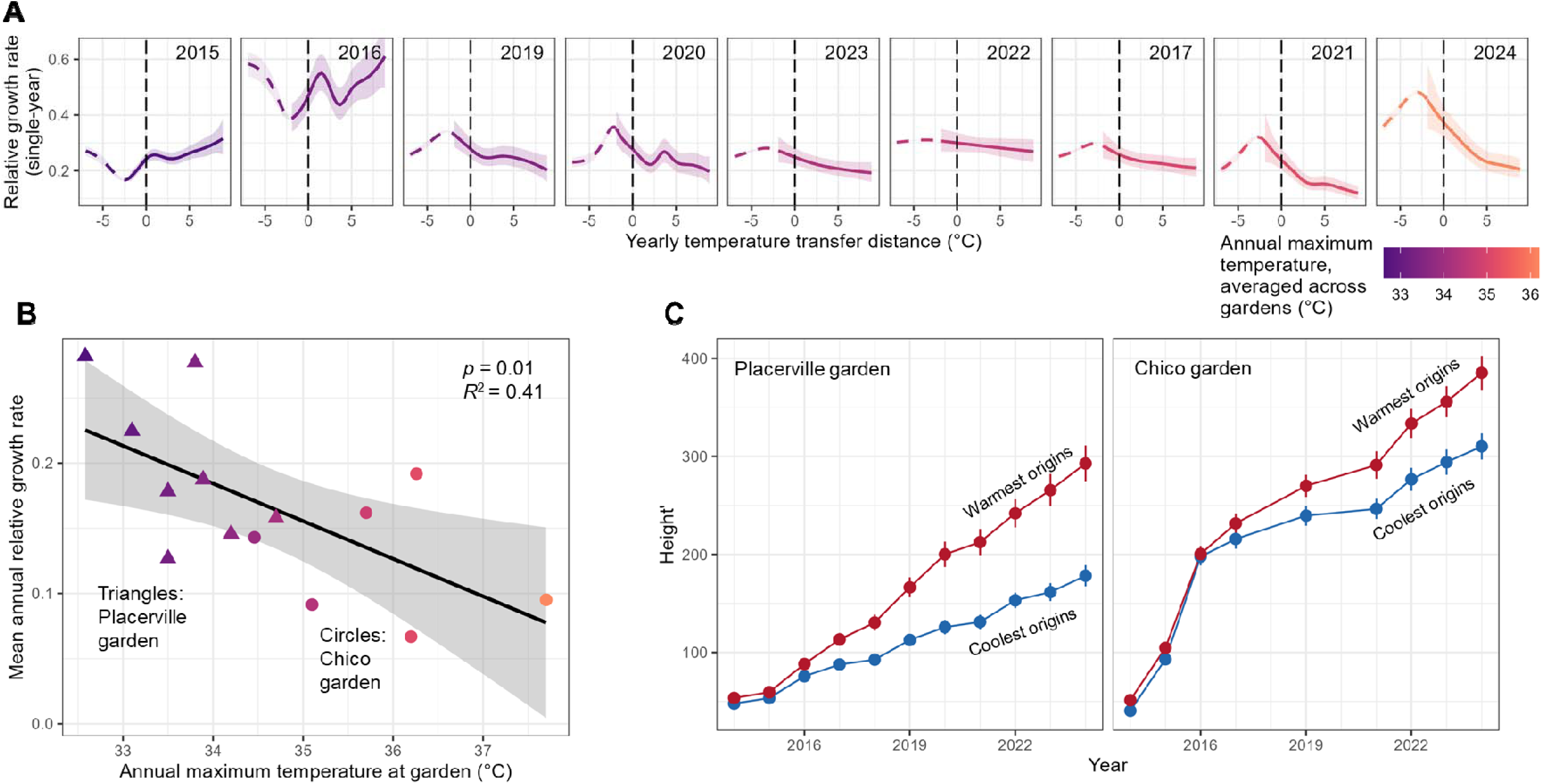
Exacerbated patterns of maladaptation in warm years lead to a lasting performance gap among valley oak progeny in a common garden. A: Strength of relationship between temperature transfer distance and relative growth rate significantly differs among study years (*p*_Year×Tdiffyear_ > 0.001 in all years). Solid colored curves represent empirically derived generalized additive model predictions of single-year relative growth rate as a function of yearly temperature transfer distance across the period of tree growth in the common gardens, 2015-2024 (2018 is excluded due to incomplete data). Long-dashed curves represent extrapolations of the model using Gaussian process regression. Vertical dashed line in each plot denotes where source temperature equals garden temperature. The area to the left of the zero line contains individuals from warmer historical climates than the garden, while the area to the right contains individuals from cooler historical climates. Color of curve represents annual maximum temperature averaged across gardens. Refer to Table S10 for statistical summary of model. **B:** Average relative growth rates are lower in hotter years. Each point represents the average relative growth rate of all trees in one garden in one year. Years 2015 and 2016 are not shown here because all trees had exceptionally high growth in those years, possibly due to maternal or outplanting effects. Color represents the annual maximum temperature at the garden. Black line is the best-fit line ± 95% confidence interval from a linear regression (n = 14). **C:** Performance gap between trees from warm and cool origins planted in two common gardens increases over time in both common gardens, especially in the hottest study years. Refer to Table S11 for summary of the ANCOVA represented by this figure. For aid in visualizing patterns over time, tree height′ (see Eq. S1) is plotted instead of relative growth rate and only averages of families from the hottest (red lines) and coolest (blue lines) 10% of origins are shown.

### Simulated outcomes of approaches to increase future resilience

We projected a 10-30% decline in relative fitness of valley oak by 2100 with no movement of genotypes (Scenario 1, Figure 4A). The approach solely based on matching to future climate (Scenario 2) led to fitness increases in some parts of the range, but similar or even slightly worse outcomes than retaining local genotypes in other areas (Figure 4B). By contrast, the simulated performance-based future climate matching approach (Scenario 3) led to relative fitness increases up to 20% across much of the species’ range and up to 60% in several areas (Figure 4C), though there were some declines in the low-elevation central region of the range even with the performance-based approach (Figure 4C, yellow areas). (See Table S15 for statistical summary of the environmental model used to estimate the baseline distribution of relative fitness and Table S16 for statistical summary of the model used to predict future relative fitness.)

**Figure 4:**
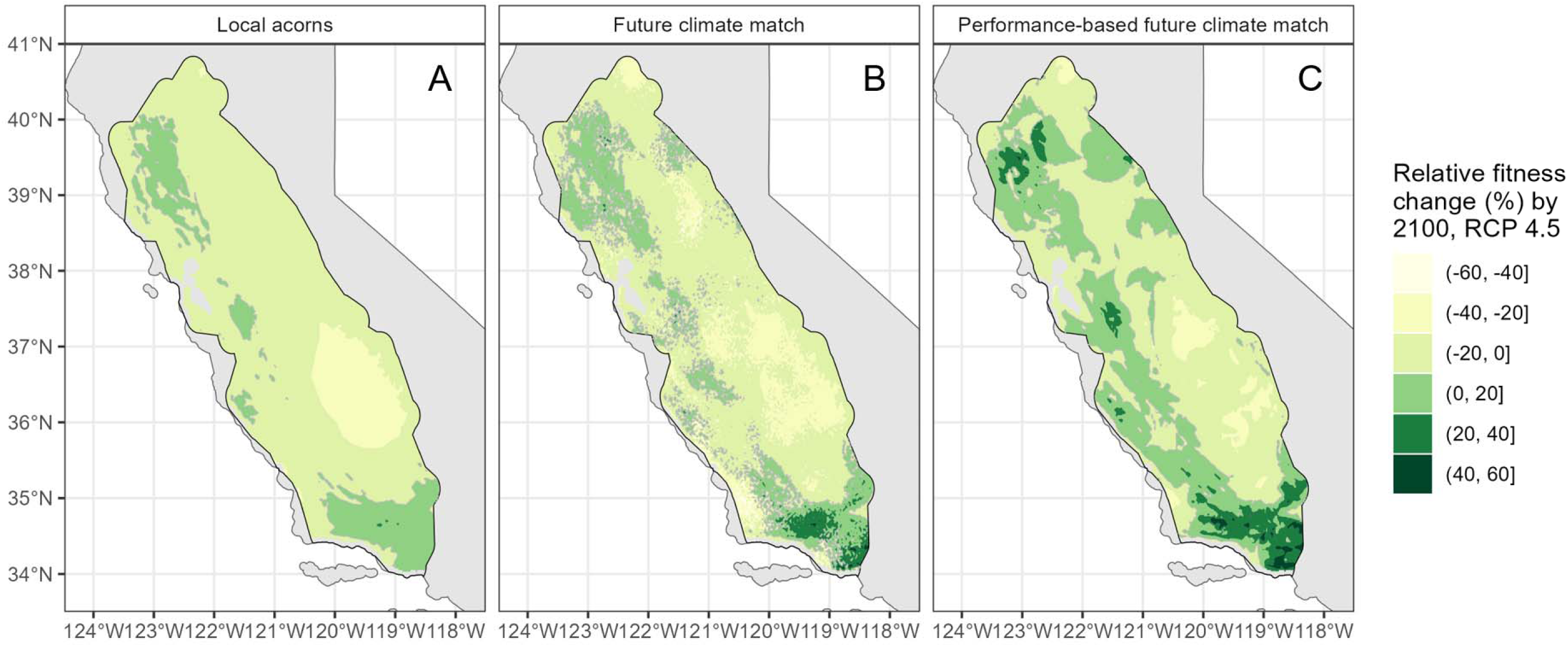
Selecting high-performing trees outperforms climate matching for establishing resilient future forests. Maps show projections of change in valley oak relative fitness across the range for years 2070-2100 under an RCP 4.5 stabilized emissions scenario. Color represents the percentage change in predicted relative fitness from present to future based on relative fitness data from the common gardens. Grey contour line shows where the change in fitness equals zero; darker areas within the line showed predicted growth increases, while lighter areas outside the line showed predicted growth decreases. The model is based on change in maximum summer temperature predicted by an ensemble of 8 global climate models downscaled for North America (Mahony *et al*. 2022). (**A**) Local acorns. (**B**) Random selection of trees from warm sites within a 25-km radius. (**C**) Selection of highest-performing trees from warm sites within a 25-km radius.

## Discussion

Adaptational lag in valley oak trees has substantially reduced their fitness, and we project a 10-30% decline in relative fitness (growth × survival) by 2100. This decline raises concern about the long-term health of oak populations in California. Importantly, amplified maladaptation in hotter years indicates that these effects are likely to intensify as temperatures rise, and that the fitness cost of stressful years can persist into subsequent growing seasons. Nonetheless, our results suggest that standing variation among valley oak families can be leveraged to improve future forest resilience. Identifying high-performing phenotypes has the potential to improve population-level outcomes, which can help conserve valley oak populations by reducing ongoing maladaptation.

### Persistent maladaptation to current temperature portends vulnerability to climate change

Valley oak trees at 12 years exhibit maladaptation to contemporary summer temperatures as we observed for five-year old trees (Browne *et al*. 2019), suggesting a persistent adaptational lag to climates dating back to the Last Glacial Maximum. Previous studies have mostly found evidence of current maladaptation in young trees up to 4 years old (Gellie *et al*. 2016; Butnor *et al*. 2019; Etterson *et al*. 2020). Here, we observed a persistent maladaptive pattern of growth across a 12-year longitudinal study, which provides robust evidence that maladaptation persists across the juvenile stage rather than diminishing with age. Because anthropogenic climate change is occurring much more quickly than any previous natural climate change (Intergovernmental Panel on Climate Change 2023), valley oaks’ inability to keep pace with past climate change raises serious concern about their future resilience.

### Contrasting adaptation to winter and summer temperatures

In contrast to summer temperatures, valley oak populations appear to be locally adapted to winter temperatures. Trees from origins with similar winter temperatures to the common gardens achieved the highest growth and relative fitness, with declines observed as winter transfer distance increased. This contrast may reflect strong selective pressure imposed by freezing temperatures, which has been shown in other oak species (Bert *et al*. 2020; Ramírez-Valiente *et al*. 2022) and is supported by the prevalence of freezing winter temperatures across most of the valley oak range between the Last Glacial Maximum and the present (Karger *et al*. 2023). By contrast, summer temperature maxima (which reached 30°C in the hottest extents of the range, but were below 25°C in the rest of the range, during the Last Glacial Maximum; Karger *et al*. 2023) may have been too low to impose strong selection. After the Last Glacial Maximum, summer temperatures increased more rapidly than winter temperatures (Karger *et al*. 2023), potentially allowing winter adaptation to keep pace while summer adaptation lagged behind. However, despite apparent local adaptation to winter temperatures, adaptational lag to summer temperature represents a primary vulnerability for valley oak populations under future warming due to its strong association with reduced growth and fitness.

### Maladaptation is amplified in hotter years

Exacerbated maladaptation in hotter years demonstrated that valley oaks are not only maladapted to average temperatures but also sensitive to short-term temperature extremes. While warm-origin trees outperformed cool-origin trees in hot years, all trees performed worse on average under higher temperatures regardless of origin, with no indication of an optimum temperature or range. Notably, the cumulative temperature-fitness relationships show evidence of steeper declines in past fitness than in predicted future fitness, suggesting that future declines may be minimal. However, we caution against this interpretation because interannual variation in the severity of the climate-fitness relationship demonstrates that acute stresses from warm years can greatly exacerbate the effects of adaptational lag. In addition, while past changes in temperature and fitness occurred over roughly 20,000 years, future changes of similar magnitude will occur within the lifetimes of currently living trees (AdaptWest 2022). We therefore find that valley oaks are vulnerable to increasing average temperatures as well as increasingly frequent and severe extreme weather events, as has been observed in other species (Benito Garzón *et al*. 2013a; Frank *et al*. 2017a; Derry *et al*. 2019).

This vulnerability indicates limited capacity for physiological buffering, acclimation, or adaptive plasticity to offset increasing thermal stress. This result contrasts with findings of adaptive plasticity and acclimation in other oak species in response to temperature (Bert *et al*. 2020; Martínez-Sancho *et al*. 2025), suggesting that valley oak is especially vulnerable to climate change. Gauthey *et al*. (2024) found that three Mediterranean oak species could endure air temperature over 50°C without leaf mortality but net photosynthesis approached zero at ∼35°C, which could explain why we observed growth declines but not significant mortality in hotter years. In the hottest study year at the Chico garden, which also had the lowest average relative growth rate, maximum temperatures exceeded those experienced historically by any maternal seed source, suggesting that parts of the species range may already be approaching a temperature threshold (Normand *et al*. 2009; Beigaitė *et al*. 2022; Osland *et al*. 2025). Continued warming is therefore likely to constrain growth across the species’ range, and especially in hotter regions.

### Simulated assisted gene flow shows potential to bolster population resilience to future climate

Despite an overall pattern of climate maladaptation in valley oak, high standing variation in fitness responses among families suggests that we can improve the resilience of future forests through movement of existing genotypes. Highly outcrossing species like oaks contain some individuals that are well-adapted to the local conditions and others that are less adapted (Sork *et al*. 2013; Brady *et al*. 2019a). The results of our simulations indicate potential for high-performing genotypes “preadapted” to future conditions (*sensu* Aitken & Bemmels 2016) to be identified throughout the species range and that planting these preadapted genotypes can result in more resilient future populations than an approach guided only by climate matching alone. Thus, matching purely by climate without considering growth variation risks selecting trees that are already maladapted to their current environments and would thus be equally maladapted to the matched future climates in which they are planted. High phenotypic variation among families—even from similar climates—has been observed in other provenance tests (Wang *et al*. 2010; Benito Garzón *et al*. 2019), suggesting that a performance-based climate matching strategy may be applicable to other species if they have sufficient individual phenotypic variation (e.g., Beaulieu & Rainville 2005; Robert *et al*. 2024).

Our findings also support a key underlying assumption of landscape genomic approaches: That adaptedness should be related to phenotypic performance (e.g., Mead *et al*. 2024; Muniz *et al*. 2024; Buck *et al*. 2026a; Buck *et al*. 2026b). Validating conservation genomic methods with empirical data is a current area of urgency (Archambeau *et al*. 2024; Lotterhos 2024b; Fitzpatrick *et al*. 2026). By quantifying growth and survival, we provide empirical calibration of genomic forecasts and a framework for interpreting the genomic adaptedness index in terms of realized fitness consequences. Such comparisons can aid in genomic forecasting by calibrating models to account for existing maladaptation (i.e., training them on historical climate data), thus avoiding fundamental limitations caused by the assumption that populations are locally adapted to current climates. Furthermore, our findings in 12-year-old trees amplify the conclusions of Browne et al. (2019) noted in 5-year-old seedlings and therefore illustrate that it is possible to validate genome-informed recommendations using growth patterns of young trees, which is valuable because it will not always be feasible to conduct long-term (10–20+ years) common garden studies due to limited resources or urgency in making management decisions (Benito Garzón *et al*. 2013b; Bowman *et al*. 2013; Huxman *et al*. 2022). More studies are needed to directly compare phenotypic and genomic predictions of response to climate change (but see Archambeau *et al*. 2024; Verrico *et al*. 2026) and work is in progress on this study system towards that end. Integrating phenotypic and genomic information will be essential for improving predictions and guiding conservation decisions under rapid climate change.

### Implications for management

Our findings from a long-term, range-wide common garden study can inform seed sourcing strategies for *Quercus lobata* and suggest that similar methods can inform conservation of other species. The results of our simulations suggest that high-performing trees (i.e., trees that grow well in warmer climates) can be identified throughout the species’ range despite a general pattern of maladaptation to climate. Thus, this approach can substantially bolster resilience of the species by augmenting populations with seeds from high-performing trees growing in warmer localities than the planting site. By contrast, not incorporating climate data risks fitness declines across most of the species’ range.

Common garden experiments provide a powerful tool for management by identifying which trees from a locality are likely to perform well at a given planting site. To illustrate this point, consider a restoration project that seeks to establish additional oak trees at a site. Based on the predicted future climate of that site, we would first identify localities whose current climates match the projected future climate of the planting site based on climate transfer functions. We would then determine whether trees within those localities include trees that grow well in our common garden. Finally, we could visit these climate-matched localities and prioritize acorn collection from trees identified as high performers in the common gardens. However, limiting restoration projects to using seeds from only these confirmed high performers would limit the potential genetic diversity of restored populations (Vander Mijnsbrugge *et al*. 2010; Byrne *et al*. 2011; Derry *et al*. 2019). Thus, we could selectively collect seeds from the best performing seed producers at that locality, thereby increasing the likelihood of capturing relatively high-performing individuals. Acknowledging that those good seed producers may be growing well for non-genetic reasons, they are still better candidates for restoration projects than poor-growing trees. Indeed, many studies have found a relationship between tree size and seed production (Younginger *et al*. 2017). This approach would not be expected to identify high performers with the same rigor as a long-term common garden study (Huxman *et al*. 2022), but we would expect an outcome somewhere between a purely climate-matching strategy and an ideal performance-informed climate-matching strategy. Because the potential negative effects of leaving genotypes in place are widespread, even imperfect strategies that account for maladaptation to climate change may improve outcomes relative to approaches that do not.

Common garden experiments can also be combined with genomic information to identify genotypes with faster growth rates under warmer conditions. Simulations by Browne *et al*. (2019) suggest that incorporating such genomic information into seed sourcing could substantially increase growth under future climates. Restoration projects that have access to common garden and genomic data could screen climate-matched localities for individuals with high numbers of these performance-associated alleles. As genomic resources become increasingly accessible, managers could ultimately improve the precision and scalability of assisted gene flow by integrating phenotypic, genomic, and climate data to identify genetic variation associated with high performance under future climates without requiring long-term common garden evaluation of every potential seed source. In the meantime, we suggest that small- or large-scale common garden experiments can be valuable resources for informing management decisions.

### Caveats

Our inferences must be considered within the context of the assumptions of the study. First, we observed two components of fitness of individuals (growth and survival), but not demographic rates of populations because we do not yet have fecundity data, which is a common limitation in studies of long-lived species (Bowman *et al*. 2013; Aspalter *et al*. 2025). We are also inherently limited in our ability to extrapolate patterns of relative fitness in 12-year-old juvenile trees to adult trees that can live hundreds of years. Provenance tests of other species have shown that signals of adaptation can change drastically over time (e.g., Bennington *et al*. 2012; Germino *et al*. 2019), although others have found these signals to remain highly consistent (e.g., Kashian & Barnes 2021; Chmura & Modrzyński 2023). Thus, our findings should not be interpreted as representative of mature trees’ lifetime fitness.

Second, our models and thus our future predictions are only based on changes in maximum summer temperatures. While we, and others (e.g., Ramírez-Valiente *et al*. 2010; Ramírez-Valiente *et al*. 2022), have found maximum temperature to be a predominant selective pressure shaping climate adaptation of oak trees in Mediterranean climates, future climate change is predicted to alter multiple climate variables that may also exert selective pressure (Anderson 2016; Solé-Medina *et al*. 2022). Water availability is commonly predicted to affect future plant adaptedness, especially in California (e.g., McLaughlin & Zavaleta 2012; Mead *et al*. 2019), but our ability to assess the effects of local precipitation on growth in the gardens was limited because they were irrigated. Regardless, precipitation was strongly inversely correlated with summer maximum temperature at the origin sites, which suggests that adaptation to drought covaries with adaptation to high temperatures. Furthermore, water availability varies at the microclimate level due to soil and hydrology, so predictions based only on precipitation may not accurately reflect the stresses experienced by plants (Dekker *et al*. 2007; Fan *et al*. 2017).

Third, another potentially important climate characteristic is that while mean winter temperature is predicted to increase by 2100 under the RCP 4.5 scenario, freezes will still occur (AdaptWest 2022) and anomalous late freezing events may become more frequent (Benito Garzón *et al*. 2013a; Wang *et al*. 2017). Therefore, conservation planning that only seeks to maximize adaptedness to hot extremes may result in populations that are maladapted to cold temperatures. However, negative effects of rapidly increasing temperatures on plants are well documented (Parmesan & Hanley 2015), so our inferences are biologically justified.

### Conclusions

Maladaptation to contemporary and future climates is already constraining the performance of a long-lived keystone tree species in California. Using a 12-year, range-wide provenance test, we show a climate–fitness mismatch that intensifies during hotter years, imposes lasting fitness penalties, and does not diminish with age, demonstrating sustained adaptational lag rather than transient maternal effects or acclimation.

Despite pervasive maladaptation, substantial genetically based phenotypic variation remains among families. By exploiting this standing variation, performance-based climate matching can yield greater projected future performance than climate matching alone, however climate matching can still provide an improvement. Together, our results provide novel phenotypic evidence that maladaptation is already operating at ecologically meaningful scales, that climate variability amplifies its fitness costs, and that explicitly accounting for current maladaptation shows promise for managing keystone species under rapid climate change.

Statement of authorship

VLS and JWW designed the common garden experiment; VLS, JWW, and ARBG designed research questions; all authors performed research; MO and BB contributed additional data; ARBG analyzed data with input from VLS; ARBG and VLS wrote the paper with input from JWW.

## Data availability statement

Data and scripts underlying the analyses in this paper are deposited in FigShare (10.6084/m9.figshare.31937499). All data processing and analysis scripts are publicly available through the lead author’s personal GitHub page (github.com/arbgoetz). Finally, all raw data are publicly available through the project’s interactive data portal (sorklab2.eeb.ucla.edu/app/).

## Acknowledgements

We acknowledge the native peoples of California (past, present, and future) as the traditional stewards of California oak ecosystems. For their assistance in establishing and maintaining the provenance test, we thank Annette Delfino Mix, Courtney Canning, and Lisa Crane. We thank Paul Gugger who organized the 2012 teams, as well as the numerous people who collected the 11,000 acorns needed to start the provenance test. We also thank the many past and present Sork Lab members and volunteers who assisted with annual data collection. Finally, we thank three anonymous reviewers and the other current Sork Lab members (Ryan Buck, Lily Peck, Heidi Yang, and Jacqueline Holmes) for substantive feedback on this manuscript. This research was funded by an NSF long-term research grant awarded to VLS and JWW (LTREB-2232794), the USDA Forest Service Pacific Southwest Research Station, and UCLA.

The findings and conclusions in this publication are those of the authors and should not be construed to represent any official USDA or U.S. Government determination or policy. Any use of product names is for informational purposes only and does not imply endorsement by the US Government.

## References

AdaptWest, P. (2022). Gridded current and projected climate data for North America at 1km resolution, generated using the ClimateNA v7.30 software (T. Wang et al., 2022). Data Basin adaptwest.databasin.org.

AdaptWest Project (2022). Gridded current and projected climate data for North America at 1km resolution, generated using the ClimateNA v7.30 software (T. Wang et al., 2022). Data Basin adaptwest.databasin.org.

Aitken, S.N. & Bemmels, J.B. (2016). Time to get moving: assisted gene flow of forest trees. Evolutionary Applications, 9, 271–290.

Aitken, S.N., Yeaman, S., Holliday, J.A., Wang, T. & Curtis-McLane, S. (2008). Adaptation, migration or extirpation: climate change outcomes for tree populations. Evolutionary Applications, 1, 95–111.

Anderson, J.T. (2016). Plant fitness in a rapidly changing world. New Phytologist, 210, 81–87.

Anderson, J.T., DeMarche, M.L., Denney, D.A., Breckheimer, I., Santangelo, J. & Wadgymar, S.M. (2025). Adaptation and gene flow are insufficient to rescue a montane plant under climate change. Science, 388, 525–531.

Anderson, J.T. & Wadgymar, S.M. (2020). Climate change disrupts local adaptation and favours upslope migration. Ecology Letters, 23, 181–192.

Archambeau, J., Benito Garzón, M., de-Miguel, M., Changenet, A., Bagnoli, F., Barraquand, F., et al. (2024). Evaluating genomic offset predictions in a forest tree with high population genetic structure. Evolutionary Biology.

Ashley, M.V., Abraham, S.T., Backs, J.R. & Koenig, W.D. (2015). Landscape genetics and population structure in Valley Oak (Quercus lobata Née). American Journal of Botany, 102, 2124–2131.

Aspalter, S., Ciceu, A., Landivar Albis, C.M., Chakraborty, D. & Schueler, S. (2025). Provenance variation in functional traits of European forest trees: Meta-analysis reveals effects of taxa and age despite critical research gaps. Ecol Evol, 15, e71834.

Beaulieu, J. & Rainville, A. (2005). Adaptation to climate change: Genetic variation is both a short- and a long-term solution. The Forestry Chronicle, 81, 704–709.

Beigaitė, R., Tang, H., Bryn, A., Skarpaas, O., Stordal, F., Bjerke, J.W. et al. (2022). Identifying climate thresholds for dominant natural vegetation types at the global scale using machine learning: Average climate versus extremes. Global Change Biology, 28, 3557–3579.

Benito Garzón, M., Ha-Duong, M., Frascaria-Lacoste, N. & Fernández Manjarrés, J. (2013a). Extreme climate variability should be considered in forestry assisted migration. BioScience, 63, 317–317.

Benito Garzón, M., Ha-Duong, M., Frascaria-Lacoste, N. & Fernández Manjarrés, J. (2013b). Habitat restoration and climate change: dealing with climate variability, incomplete data, and management decisions with tree translocations. Restoration Ecology, 21, 530–536.

Benito Garzón, M., Robson, T.M. & Hampe, A. (2019). ΔTraitSDMs: species distribution models that account for local adaptation and phenotypic plasticity. New Phytologist, 222, 1757–1765.

Bennington, C.C., Fetcher, N., Vavrek, M.C., Shaver, G.R., Cummings, K.J. & McGraw, J.B. (2012). Home site advantage in two long-lived arctic plant species: results from two 30-year reciprocal transplant studies. Journal of Ecology, 100, 841–851.

Benomar, L., Bousquet, J., Perron, M., Beaulieu, J. & Lamara, M. (2022). Tree maladaptation under mid-latitude early spring warming and late cold spell: implications for assisted migration. Frontiers in Plant Science, 13.

Bert, D., Lebourgeois, F., Ponton, S., Musch, B. & Ducousso, A. (2020). Which oak provenances for the 22nd century in Western Europe? Dendroclimatology in common gardens. PLOS ONE, 15, e0234583.

Borgman, E.M., Schoettle, A.W. & Angert, A.L. (2015). Assessing the potential for maladaptation during active management of limber pine populations: a common garden study detects genetic differentiation in response to soil moisture in the Southern Rocky Mountains. Canadian Journal of Forest Research, 45, 496–505.

Bowman, D.M.J.S., Brienen, R.J.W., Gloor, E., Phillips, O.L. & Prior, L.D. (2013). Detecting trends in tree growth: not so simple. Trends in Plant Science, 18, 11–17.

Brady, S.P., Bolnick, D.I., Angert, A.L., Gonzalez, A., Barrett, R.D.H., Crispo, E. et al. (2019a). Causes of maladaptation. Evolutionary Applications, 12, 1229–1242.

Brady, S.P., Bolnick, D.I., Barrett, R.D.H., Chapman, L., Crispo, E., Derry, A.M. et al. (2019b). Understanding maladaptation by uniting ecological and evolutionary perspectives. The American Naturalist, 194, 495–515.

Browne, L., Wright, J.W., Fitz-Gibbon, S., Gugger, P.F. & Sork, V.L. (2019). Adaptational lag to temperature in valley oak (*Quercus lobata*) can be mitigated by genome-informed assisted gene flow. Proceedings of the National Academy of Sciences of the United States of America, 116, 25179–25185.

Buck, R.C., Butterfield, H.S., Hiroyasu, E., Howard, J., Knapp, J., Principe, Z., et al. (2026a). Landscape genomic analysis of *Quercus agrifolia* Née predicts patterns of adaptedness to future climate and provides guidance for conservation. Evolutionary Applications, In Press.

Buck, R.C., Butterfield, H.S., Hiroyasu, E.H.T., Howard, J., Principe, Z., Rose, M.B. et al. (2026b). Applying both landscape genomic and ecological niche model predictions to inform conservation strategies of a California foundational oak species. Molecular Ecology, 35, e70322.

Butnor, J.R., Verrico, B.M., Johnsen, K.H., Maier, C.A., Vankus, V. & Keller, S.R. (2019). Phenotypic variation in climate-associated traits of red spruce (*Picea rubens* Sarg.) along elevation gradients in the southern Appalachian Mountains. Castanea, 84, 128–143.

Byrne, M., Stone, L. & Millar, M.A. (2011). Assessing genetic risk in revegetation. Journal of Applied Ecology, 48, 1365–1373.

Callaham, R.Z. (1963). Provenance research: investigation of genetic diversity associated with geography. Unasylua, 18 (2-3), 2–12.

Capblancq, T., Fitzpatrick, M.C., Bay, R.A., Exposito-Alonso, M. & Keller, S.R. (2020). Genomic prediction of (mal)adaptation across current and future climatic landscapes. Annual Review of Ecology, Evolution, and Systematics, 51, 245–269.

Carter, K.K. (1996). Provenance tests as indicators of growth response to climate change in 10 north temperate tree species. Canadian Journal of Forest Research, 26, 1089–1095.

Chakraborty, D., Schueler, S., Lexer, M.J. & Wang, T. (2019). Genetic trials improve the transfer of Douglas-fir distribution models across continents. Ecography, 42, 88–101.

Chmura, D.J. & Modrzyński, J. (2023). Sensitivity of height growth response to climate change does not vary with age in common garden among Norway spruce populations from elevational gradients. Forest Ecology and Management, 542, 121118.

Close, D.C., Beadle, C.L. & Brown, P.H. (2005). The physiological basis of containerised tree seedling ‘transplant shock’: a review. Australian Forestry, 68, 112–120.

Corlett, R.T. & Westcott, D.A. (2013). Will plant movements keep up with climate change? Trends in Ecology & Evolution, 28, 482–488.

Davis, F.W., Stoms, D.M., Hollander, A.D., Thomas, K.A., Stine, P.A., Odion, D., et al. (1998). The California Gap Analysis Project: Final Report. p. 255.

Davis, M.B. & Shaw, R.G. (2001). Range shifts and adaptive responses to quaternary climate change. Science, 292, 673–679.

Davis, M.B., Shaw, R.G. & Etterson, J.R. (2005). Evolutionary responses to changing climate. Ecology, 86, 1704–1714.

Dekker, S.C., Rietkerk, M. & Bierkens, M.F.P. (2007). Coupling microscale vegetation–soil water and macroscale vegetation–precipitation feedbacks in semiarid ecosystems. Global Change Biology, 13, 671–678.

Delfino Mix, A., Wright, J.W., Gugger, P.F., Liang, C. & Sork, V.L. (2015). Establishing a range-wide provenance test in valley oak (*Quercus lobata* Née) at two California sites. US Forest Service, US Department of Agriculture, pp. 413–424.

Depardieu, C., Girardin, M.P., Nadeau, S., Lenz, P., Bousquet, J. & Isabel, N. (2020). Adaptive genetic variation to drought in a widely distributed conifer suggests a potential for increasing forest resilience in a drying climate. New Phytologist, 227, 427–439.

Derry, A.M., Fraser, D.J., Brady, S.P., Astorg, L., Lawrence, E.R., Martin, G.K. et al. (2019). Conservation through the lens of (mal)adaptation: Concepts and meta-analysis. Evolutionary Applications, 12, 1287–1304.

Diamond, S.E. & Martin, R.A. (2020). Evolution is a double-edged sword, not a silver bullet, to confront global change. Annals of the New York Academy of Sciences, 1469, 38–51.

Dutech, C., Sork, V.L., Irwin, A.J., Smouse, P.E. & Davis, F.W. (2005). Gene flow and fine-scale genetic structure in a wind-pollinated tree species, *Quercus lobata* (Fagaceaee). American Journal of Botany, 92, 252–261.

Ellison, A.M. (2019). Foundation species, non-trophic interactions, and the value of being common. iScience, 13, 254–268.

Erickson, C. (2024). GauPro: Gaussian Process Fitting.

Etterson, J.R., Cornett, M.W., White, M.A. & Kavajecz, L.C. (2020). Assisted migration across fixed seed zones detects adaptation lags in two major North American tree species. Ecological Applications, 30, e02092.

Falconer, D.S. & Mackay, T.F. (1983). Quantitative Genetics. Longman London.

Fan, Y., Miguez-Macho, G., Jobbágy, E.G., Jackson, R.B. & Otero-Casal, C. (2017). Hydrologic regulation of plant rooting depth. Proceedings of the National Academy of Sciences, 114, 10572–10577.

Fitzpatrick, M.C., Keller, S.R. & Lotterhos, K.E. (2026). The Challenge of Genomic Forecasting in an Era of Global Change. The American Naturalist, 000–000.

Flint, L.E., Flint, A.L., Thorne, J.H. & Boynton, R. (2013). Fine-scale hydrologic modeling for regional landscape applications: the California Basin Characterization Model development and performance. Ecological Processes, 2, 25.

Foest, J.J., Szymkowiak, J., Dyderski, M.K., Jastrzębowski, S., Fuchs, H., Ratajczak, E. et al. (2025). No refuge at the edge for European beech as climate warming disproportionately reduces masting at colder margins. Ecology Letters, 28, e70284.

Frank, A., Howe, G.T., Sperisen, C., Brang, P., St. Clair, J.B., Schmatz, D.R. et al. (2017a). Risk of genetic maladaptation due to climate change in three major European tree species. Global Change Biology, 23, 5358–5371.

Frank, A., Sperisen, C., Howe, G.T., Brang, P., Walthert, L., St.Clair, J.B., et al. (2017b). Distinct genecological patterns in seedlings of Norway spruce and silver fir from a mountainous landscape. Ecology, 98, 211–227.

Gauthey, A., Kahmen, A., Limousin, J.M., Vilagrosa, A., Didion-Gency, M., Mas, E. et al. (2024). High heat tolerance, evaporative cooling, and stomatal decoupling regulate canopy temperature and their safety margins in three European oak species. Glob Chang Biol, 30, e17439.

Gellie, N.J.C., Breed, M.F., Thurgate, N., Kennedy, S.A. & Lowe, A.J. (2016). Local maladaptation in a foundation tree species: Implications for restoration. Biological Conservation, 203, 226–232.

Germino, M.J., Moser, A.M. & Sands, A.R. (2019). Adaptive variation, including local adaptation, requires decades to become evident in common gardens. Ecological Applications, 29, e01842.

Girardin, M.P., Hogg, E.H., Bernier, P.Y., Kurz, W.A., Guo, X.J. & Cyr, G. (2016). Negative impacts of high temperatures on growth of black spruce forests intensify with the anticipated climate warming. Global Change Biology, 22, 627–643.

Girardin, M.P., Isabel, N., Guo, X.J., Lamothe, M., Duchesne, I. & Lenz, P. (2021). Annual aboveground carbon uptake enhancements from assisted gene flow in boreal black spruce forests are not long-lasting. Nature Communications, 12, 1169.

Gougherty, A.V., Keller, S.R. & Fitzpatrick, M.C. (2021). Maladaptation, migration and extirpation fuel climate change risk in a forest tree species. Nat. Clim. Chang., 11, 166–171.

Gugger, P.F., Ikegami, M. & Sork, V.L. (2013). Influence of late Quaternary climate change on present patterns of genetic variation in valley oak, *Quercus lobata* Née. Molecular Ecology, 22, 3598–3612.

Hastie, T. & Tibshirani, R. (1986). Generalized Additive Models. Statistical Science, 1, 297–310.

Hendry, A.P. & Gonzalez, A. (2008). Whither adaptation? Biology & Philosophy, 23, 673–699.

Huxman, T.E., Winkler, D.E. & Mooney, K.A. (2022). A common garden super-experiment: An impossible dream to inspire possible synthesis. Journal of Ecology, 110, 997–1004.

Intergovernmental Panel on Climate Change (2023). Climate Change 2021 – The Physical Science Basis: Working Group I Contribution to the Sixth Assessment Report of the Intergovernmental Panel on Climate Change. 1 edn. Cambridge University Press.

Jump, A.S. & Peñuelas, J. (2005). Running to stand still: adaptation and the response of plants to rapid climate change. Ecology Letters, 8, 1010–1020.

Karger, D.N. (2025). CHELSA-TraCE21k-centennial-bioclim and topographic data since the Last Glacial Maximum. EnviDat.

Karger, D.N., Conrad, O., Böhner, J., Kawohl, T., Kreft, H., Soria-Auza, R.W. et al. (2017). Climatologies at high resolution for the earth’s land surface areas. Sci Data, 4, 170122.

Karger, D.N., Nobis, M.P., Normand, S., Graham, C.H. & Zimmermann, N.E. (2023). CHELSA-TraCE21k – high-resolution (1-km) downscaled transient temperature and precipitation data since the Last Glacial Maximum. Climate of the Past, 19, 439–456.

Kashian, D.M. & Barnes, B.V. (2021). Tree growth and survival over 61 years at the Second International Larch Provenance Test in southeastern Michigan, USA. Silvae Genetica, 70, 9–21.

Kelly, P.F., Phillips, S.E. & Williams, D.F. (2005). Documenting ecological change in time and space: The San Joaquin Valley of California. In: Mammalian Diversification: From Chromosomes to Phylogeography (A Celebration of the Career of James L. Patton (eds. Lacey, EA & Myers, P). University of California Press Berkeley and Los Angeles, CA.

Kooyers, N.J., Anderson, J.T., Angert, A.L., Avolio, M.L., Campbell, D.R., Exposito-Alonso, M. et al. (2025). Responses to climate change – insights and limitations from herbaceous plant model species. New Phytologist, 248, 461–493.

Kueppers, L.M., Snyder, M.A., Sloan, L.C., Zavaleta, E.S. & Fulfrost, B. (2005). Modeled regional climate change and California endemic oak ranges. Proceedings of the National Academy of Sciences, 102, 16281–16286.

LaDochy, S. & Witiw, M. (2023). Climate Change in California: Past, Present, and Future. In: *Fire and Rain*, pp. 163–183.

Leites, L. & Benito Garzón, M. (2023). Forest tree species adaptation to climate across biomes: Building on the legacy of ecological genetics to anticipate responses to climate change. Global Change Biology, 29, 4711–4730.

Lind, B.M., Candido-Ribeiro, R., Singh, P., Lu, M., Obreht Vidakovic, D., Booker, T.R. et al. (2024). How useful is genomic data for predicting maladaptation to future climate? Global Change Biology, 30, e17227.

Lloret, F., Keeling, E.G. & Sala, A. (2011). Components of tree resilience: effects of successive low-growth episodes in old ponderosa pine forests. Oikos, 120, 1909–1920.

Lotterhos, K.E. (2024a). Interpretation issues with “genomic vulnerability” arise from conceptual issues in local adaptation and maladaptation. Evolution Letters, 8, 331–339.

Lotterhos, K.E. (2024b). Principles in experimental design for evaluating genomic forecasts. Methods in Ecology and Evolution, 15, 1466–1482.

Mahony, C.R., Wang, T., Hamann, A. & Cannon, A.J. (2022). A global climate model ensemble for downscaled monthly climate normals over North America. International Journal of Climatology, 42, 5871–5891.

Martínez-Sancho, E., Rellstab, C., Fonti, P., Benito Garzón, M., Bigler, C., Miranda, J.C. et al. (2025). Genetic and plastic effects on trait variability in two major tree species: Insights from common garden experiments across Europe. Forest Ecology and Management, 597, 123126.

Mátyás, C. (1996). Climatic adaptation of trees: rediscovering provenance tests. Euphytica, 92, 45–54.

McKay, J.K., Christian, C.E., Harrison, S. & Rice, K.J. (2005). “How local is local?”—A review of practical and conceptual issues in the genetics of restoration. Restoration Ecology, 13, 432–440.

McLaughlin, B. & Zavaleta, E.S. (2012). Predicting species responses to climate change: demography and climate microrefugia in California valley oak (*Quercus lobata*). Global Change Biology, 18, 2301–2312.

Mead, A., Fitz-Gibbon, S., Knapp, J. & Sork, V.L. (2024). Comparison of conservation strategies for California Channel Island Oak (*Quercus tomentella*) using climate suitability predicted from genomic data. Evolutionary Applications, 20, 1–20.

Mead, A., Peñaloza Ramirez, J., Bartlett, M.K., Wright, J.W., Sack, L. & Sork, V.L. (2019). Seedling response to water stress in valley oak (*Quercus lobata)* is shaped by different gene networks across populations. Molecular Ecology, 28, 5248–5264.

Merlin, M., Duputié, A. & Chuine, I. (2018). Limited validation of forecasted northward range shift in ten European tree species from a common garden experiment. Forest Ecology and Management, 410, 144–156.

Muniz, A.C., de Lemos, J.P. & Lovato, M.B. (2024). Non-adaptedness and vulnerability to climate change threaten *Plathymenia* trees (Fabaceae) from the Cerrado and Atlantic Forest. Scientific Reports, 14.

Normand, S., Treier, U.A., Randin, C., Vittoz, P., Guisan, A. & Svenning, J.-C. (2009). Importance of abiotic stress as a range-limit determinant for European plants: insights from species responses to climatic gradients. Global Ecology and Biogeography, 18, 437–449.

O’Neill, G.A., Stoehr, M. & Jaquish, B. (2014). Quantifying safe seed transfer distance and impacts of tree breeding on adaptation. Forest Ecology and Management, 328, 122–130.

Osland, M.J., Bradford, J.B., Toth, L.T., Germino, M.J., Grace, J.B., Drexler, J.Z. et al. (2025). Ecological thresholds and transformations due to climate change: The role of abiotic stress. Ecosphere, 16, e70229.

Parmesan, C. & Hanley, M.E. (2015). Plants and climate change: complexities and surprises. Annals of Botany, 116, 849–864.

Pavlik, B.M., Muick, P.C., Johnson, S.G. & Popper, M. (1991). Oaks of California. Cachuma Press, Los Olivos, CA.

Pedersen, E.J., Miller, D.L., Simpson, G.L. & Ross, N. (2019). Hierarchical generalized additive models in ecology: an introduction with mgcv. PeerJ, 7, e6876.

Ramírez-Valiente, J.A., Sánchez-Gómez, D., Aranda, I. & Valladares, F. (2010). Phenotypic plasticity and local adaptation in leaf ecophysiological traits of 13 contrasting cork oak populations under different water availabilities. Tree Physiology, 30, 618–627.

Ramírez-Valiente, J.A., Santos del Blanco, L., Alía, R., Robledo-Arnuncio, J.J. & Climent, J. (2022). Adaptation of Mediterranean forest species to climate: Lessons from common garden experiments. Journal of Ecology, 110, 1022–1042.

Rasmussen, C.E. & Williams, C.K.I. (2006). Gaussian Processes for Machine Learning. MIT Press, Cambridge, MA.

Rees, M., Osborne, Colin P., Woodward, F.I., Hulme, Stephen P., Turnbull, Lindsay A. & Taylor, Samuel H. (2010). Partitioning the components of relative growth rate: How important is plant size variation? The American Naturalist, 176, E152–E161.

Rehfeldt, G.E., Wykoff, W.R. & Ying, C.C. (2001). Physiologic plasticity, evolution, and impacts of a changing climate on *Pinus contorta*. Climatic Change, 50, 355–376.

Rellstab, C. (2021). Genomics helps to predict maladaptation to climate change. Nat. Clim. Chang., 11, 85–86.

Roach, D.A. & Wulff, R.D. (1987). Maternal effects in plants. Annual Review of Ecology and Systematics, 18, 209–235.

Robert, E., Lenz, P., Bergeron, Y., de Lafontaine, G., Bouriaud, O., Isabel, N., et al. (2024). Future carbon sequestration potential in a widespread transcontinental boreal tree species: Standing genetic variation matters! Global Change Biology, 30, e17347.

Savolainen, O., Bokma, F., Garcı a-Gil, R., Komulainen, P. & Repo, T. (2004). Genetic variation in cessation of growth and frost hardiness and consequences for adaptation of *Pinus sylvestris* to climatic changes. Forest Ecology and Management, 197, 79–89.

Savolainen, O., Pyhäjärvi, T. & Knürr, T. (2007). Gene flow and local adaptation in trees. Annual Review of Ecology, Evolution, and Systematics, 38, 595–619.

Schmeddes, J., Muffler, L., Barbeta, A., Beil, I., Bolte, A., Holm, S. et al. (2024). High phenotypic variation found within the offspring of each mother tree in *Fagus* sylvatica regardless of the environment or source population. Global Ecology and Biogeography, 33, 470–481.

Solé-Medina, A., Robledo-Arnuncio, J.J. & Ramírez-Valiente, J.A. (2022). Multi-trait genetic variation in resource-use strategies and phenotypic plasticity correlates with local climate across the range of a Mediterranean oak (*Quercus faginea*). New Phytologist, 234, 462–478.

Sork, V.L., Aitken, S.N., Dyer, R.J., Eckert, A.J., Legendre, P. & Neale, D.B. (2013). Putting the landscape into the genomics of trees: approaches for understanding local adaptation and population responses to changing climate. Tree Genetics & Genomes, 9, 901–911.

Sork, V.L., Davis, F.W., Westfall, R., Flint, A., Ikegami, M., Wang, H. et al. (2010). Gene movement and genetic association with regional climate gradients in California valley oak (*Quercus lobata* Née) in the face of climate change. Molecular Ecology, 19, 3806–3823.

St. Clair, J.B. & Howe, G.T. (2007). Genetic maladaptation of coastal Douglas-fir seedlings to future climates. Global Change Biology, 13, 1441–1454.

Thuiller, W., Albert, C., Araújo, M.B., Berry, P.M., Cabeza, M., Guisan, A. et al. (2008). Predicting global change impacts on plant species’ distributions: Future challenges. Perspectives in Plant Ecology, Evolution and Systematics, 9, 137–152.

Tyler, Claudia M., Kuhn, B. & Davis, Frank W. (2006). Demography and recruitment limitations of three oak species in California. The Quarterly Review of Biology, 81, 127–152.

Valiente-Banuet, A., Aizen, M.A., Alcántara, J.M., Arroyo, J., Cocucci, A., Galetti, M. et al. (2015). Beyond species loss: the extinction of ecological interactions in a changing world. Functional Ecology, 29, 299–307.

Vander Mijnsbrugge, K., Bischoff, A. & Smith, B. (2010). A question of origin: Where and how to collect seed for ecological restoration. Basic and Applied Ecology, 11, 300–311.

Verrico, B.M., Capblancq, T., Fitzpatrick, M.C. & Keller, S.R. (2026). Reciprocal evaluation of genomic offset predictions of climate maladaptation with independent empirical datasets. The American Naturalist, 000–000.

Wang, T., O’Neill, G.A. & Aitken, S.N. (2010). Integrating environmental and genetic effects to predict responses of tree populations to climate. Ecological Applications, 20, 153–163.

Wang, X., Jiang, D. & Lang, X. (2017). Future extreme climate changes linked to global warming intensity. Science Bulletin, 62, 1673–1680.

White, T.L. (1987). A conceptual framework for tree improvement programs. New Forests, 1, 325–342.

Wolkovich, E.M. & Donahue, M.J. (2021). How phenological tracking shapes species and communities in non-stationary environments. Biological Reviews, 96, 2810–2827.

Wood, S.N., Goude, Y. & Shaw, S. (2015). Generalized additive models for large data sets. Journal of the Royal Statistical Society Series C: Applied Statistics, 64, 139–155.

Xu, W. & Prescott, C.E. (2024). Can assisted migration mitigate climate-change impacts on forests? Forest Ecology and Management, 556, 121738.

Younginger, B.S., Sirova, D., Cruzan, M.B. & Ballhorn, D.J. (2017). Is biomass a reliable estimate of plant fitness? Appl Plant Sci, 5.

Zavaleta, E.S., Hulvey, K.B. & Fulfrost, B. (2007). Regional patterns of recruitment success and failure in two endemic California oaks. Diversity and Distributions, 13, 735–745.

