## Supplementary information for "Persistent climate maladaptation decreases performance of a keystone tree species (*Quercus lobata*)"

Supporting information for: Persistent climate maladaptation decreases performance of a keystone tree species (*Quercus lobata*)

Alexander R. B. Goetz,^1^ Marissa E. Ochoa,^1^ Berenice Badillo,^1^ Jessica W. Wright,^1,2^ Victoria L. Sork^1,3^

^1^ Department of Ecology and Evolutionary Biology, University of California, Los Angeles, CA 90095

^2^ Pacific Southwest Research Station, US Forest Service, US Department of Agriculture, Placerville, CA 95667

^3^ Institute of the Environment and Sustainability, University of California, Los Angeles, CA 90095

### **Appendices**

#### **Appendix S1. Survival and growth measurements**

At the end of each growing season from 2015-2024, survival and size were measured (except for part of the Chico Garden in 2020 due to COVID-19 restrictions). Survival was assessed visually based on observable presence of living tissue. Resprouts of individuals previously reported dead were common, so survival data were corrected when such individuals were discovered. In the early years of the study, size was measured as height of the tallest stem; starting in 2019 (when trees were 7 years old), trees at least 150 cm tall had diameter at breast height (DBH) measured in addition to height. In 2021, trees removed for thinning had biomass destructively measured. These measures were used to estimate growth. As trees grew to greater than 1.5 m (Table S1), we used DBH as a measure of size and we also used it to estimate height, with destructively measured biomass data used to supplement model training and validation (see below for details). See Table S1 for a detailed list of growth variables measured in each year. In 2018 at the Chico Garden, DBH was only measured on a haphazard subset of trees, so we were unable to estimate height for those above the 4-meter threshold; we thus excluded the 2018 Chico data to avoid biases. Because trees were thinned in 2021, our longitudinal analyses did not include any trees that were deliberately removed. We also excluded all individuals recorded as killed by damage from gophers (*Geomys* sp.), as well as individuals that showed “negative” growth at any point due to either measurement error or dieback. Thus, our mortality data minimized bias from factors not associated with climate-related selective pressures. Roughly 200 individuals out of 3,415 were excluded across all years as a result.

#### **Appendix S2. Allometric growth calculations**

To allow us to calculate relative growth rates across all years of data, we fit allometric polynomial equations to the relationship between ln(basal diameter) and ln(height) of the trees destructively sampled in 2021 (Ochoa 2024), resulting in the following formula:

**Eq. S1**. $Height = e^{0.7\ln(basal diameter) + 2.81}$ (*R*^2^ = 0.69).

To fit allometric models, we first converted DBH to basal diameter using an equation fit based on the relationship between basal diameter and DBH ($basal diameter=1.42(DBH) + 27.9$; R^2^ = 0.72) measured on a haphazard subset of trees in 2018 (n = 466). This transformation was necessary because basal diameter, not DBH, was measured in the 2021 destructive harvest (see Table S1 for more information). We then used the basal diameter/height equation to predict height from DBH for trees lacking a true height measurement from 2018-2024. The R^2^ value of the relationship between predicted height (referred to hereafter as height′) and actual height of trees withheld from the training dataset (i.e., 2021 measurements of trees not destructively sampled) was 0.92. All subsequent calculations (relative growth rate and relative fitness) were thus based on actual height when present (all individuals 2013-2017; individuals under the measurement threshold in 2018-2024 (see Table S1); individuals haphazardly sampled for complete height in 2023; all individuals at the Placerville garden in 2024) and height estimated from DBH otherwise. We note some limitations to our approach: The models did not explain all variance in height, and predicted height saturated at high DBH values. Also, some trees had multiple dominant stems but only one stem per tree was measured (except in 2023), so predicted height may be underestimated in those cases. These constraints on our measurements of size and growth are unlikely to inflate the significance of our findings, so the trends we observe remain robust.

Appendix S3. Testing for genetic differences in fitness

To test for genetic differentiation in fitness among families, we conducted a two-way ANOVA on the effect of the family*garden interaction term on 2024 height′ (standardized growth metric calculated using allometric models; see “Allometric growth calculations” above). We accounted for the effect of planting block because of known environmental differences within gardens. To determine whether relative growth rate differed among families and over time, we also conducted repeated measures ANOVAs of the family*year interaction on individuals’ relative growth rates, again controlling for planting block and with individual ID included as an error term. The repeated measures design allowed us to simultaneously test differences between individuals and within the same individuals across years. Repeated measures ANOVAs were conducted separately for each garden because we were testing whether response to the environment changed over time within gardens. All ANOVAs were conducted using the aov() function in R package stats.

Appendix S4. Calculating temperature transfer distances

To compare the climates of the maternal seed sources to those of the two gardens, we calculated transfer distances (Eriksson et al. 1980, Prescher 1986) based on maximum summer temperatures and mean winter temperatures for each family in each year, taken as the difference in degrees between temperature in each garden each year and temperature in the 30-year averages of recent-past (1951-1980) maternal source climate data. Data on maternal tree source site climates from the California Basin Characterization Model (Flint et al. 2013) were used to compare common garden growth outcomes against historical climatic conditions experienced by the maternal source trees. Maternal source climate data were 30-year averages of 1950-1981 data gridded at 270m pixel resolution (downscaled using high-resolution digital elevation models). We used 1951-1980 averages for the origin climates because they are more similar to the climates in which the maternal trees (which may be up to 300 years old, or more) would have established than modern temperatures, especially because climate was more stable across California prior to the 1990s (AdaptWest 2022). We confirmed that 1951-1980 maximum summer temperature at the study sites is correlated with estimated maximum summer temperature for the 1700-1800 and 1900-2000 centuries (Figure S1; Karger *et al.* 2023); we use climatic data from the BCM in our analyses due to its high spatial resolution.

Monthly climate data for the common garden sites were taken from PRISM (prism.oregonstate.edu) at 800m pixel resolution, using the “Explorer” feature to obtain time-series climate data at the common garden point locations for years 2013-2024. See Figure S3 for precipitation trends in the gardens across the study period. Paleoclimate data were taken from the CHELSA-TraCE12k Last Glacial Maximum dataset (Karger *et al.* 2017; Karger *et al.* 2023) at 1km resolution. Projected future climate data (Relative Concentration Pathway 4.5; “stabilization” scenario) were taken from the AdaptWest 8-model ensemble projections (AdaptWest 2022; Mahony *et al.* 2022), which use methods from ClimateNA (Wang *et al.* 2016) to downscale source rasters to 270m resolution. The ensemble projections consist of weighted averages of the following climate models: ACCESS-ESM1.5, CNRM-ESM2-1, EC-Earth3, GFDL-ESM4, GISS-E2-1-G, MIROC6, MPI-ESM1.2-HR, and MRI-ESM2.0 (Mahony *et al.* 2022). Note that here, unlike Browne et al. (2019), we present the temperature transfer distance in terms of the maternal source climate rather than the planting site (e.g., “trees from warmer climates than the garden” as opposed to “trees planted into cooler environments than the origin”) because it enhanced the clarity of the discussion of our findings. Regardless, the variable’s directionality and interpretation are the same across the two papers.

Appendix S5. Calculating relative growth rates

To compare growth across trees in the common gardens from different lineages, we calculated relative growth rates (RGR) for each tree in the common gardens on both an annual and cumulative basis using the following equation:

**Eq. S2**.	$\frac{\ln{Height'}_{2}-\ln{Height'}_{1}}{t_{2}-t_{1}}$

Height′ represents the standardized height value based on allometric models of tree size variables (Eq. S1). For the cumulative metrics, t_1_ was 2014 (the year before seedlings were outplanted in the gardens) and t_2_ was 2024. For the annual metrics, t_1_ was the year before t_2_. If a tree had missing data for one year, the preceding year was treated as t_1_ (thus, t_2_ – t_1_ was always equal to either 1 or 2 in the interannual comparisons). Prior to calculating RGR, we excluded any individuals with a recorded decrease in height or estimated height between subsequent years (generally caused by dieback or differences in rounding). We also excluded individuals that had died during the study period or that had been damaged by wildlife. Relative growth rates were not standardized by block, but differences among blocks were statistically accounted for in all tests by modeling a fixed effect of block nested within garden. Finally, relative growth rates were not calculated for the 2018 or 2020 data at the Chico Garden due to missing data (see “Survival and growth data” above for details). Thus, those years were excluded from analysis.

Appendix S6. Calculating relative fitness

To explicitly compare both tree size and survival as fitness components of the maternal tree genotypes, we calculated relative fitness values (height′ * survival) for each family in each garden. Height′ represents the standardized height value based on allometric models of tree size variables (Eq. S1). Due to known differences in height across planting blocks, particularly at the Chico Garden, we calculated individual standardized heights (dividing each tree’s height by the maximum height in its planting block, which in some cases was an estimated value) in each block and averaged across progeny of each maternal tree at each garden. Relative fitness was calculated using the following equations (*i* denotes individual progeny in gardens, *j* denotes maternal tree from which all *i* progeny descended).

**Eq. S3a**. $W_{ij}={Height'}_{ij}\times{Survival}_{ij}$

**Eq. S3b**. $W_{j}=mean RF per family =\frac{\sum_{ij} W_{ij}}{\sum i}$

**Eq. S3c**. $standardized RF= \frac{W_{j}-W_{max}}{W_{max}}$

#### **Appendix S7. Generalized additive model details**

We fit GAMs using the bam() function in package ‘mgcv’ (Wood 2011) in R 4.5.0/RStudio 2025.05.0 (Posit team 2025; R Core Team 2025). To test the predictive power of the GAMs, we calculated average root-mean squared error (RMSE) via 10-fold cross-validation using functions crossv_kfold() and rmse() in package ‘modelr’ version 0.1.11 (Wickham 2023). All numerical explanatory variables were modeled using splines with a cubic regression basis (Wood 2017; Pedersen *et al.* 2019). We used Tweedie error distributions for model fitting due to the nonnormal shape of the response variable distribution (Browne *et al.* 2019). The *k* value (number of knots used in fitting) was set to 15 for variables that indicated a poor fit with the default value (*k* = 10) and the default was used for all other variables. Because smooths are penalized in model fitting, the exact value of *k* is unimportant as long as it is not too low to capture variation in the data (Wood 2017). The following terms had *k* values set to 15: Tdiff_2014-2024_ (Eq. 1b), height_year-1_ (Eq. 2).

Maternal tree ID and locality were included as random effects in the models to account for significant differences among maternal trees (see Appendix S3 for more information) as well as spatial autocorrelation within localities. For the RGR models, we also added the fixed effect of block nested within garden to account for microclimatic differences within the gardens. In the relative fitness models, only the garden term was included because values were already standardized by block (Appendix S6). We included a measurement of initial height in 2014 (individual height in the RGR models, average height of individuals within a family in the relative fitness models) to account for differences in growth prior to outplanting.

#### **Appendix S8. Effects of phenology on yearly growth**

To determine whether phenology influenced growth across years, we tested growth rate response to date of first leaf emergence, study year, and temperature transfer distance in each year. The model was fitted using data from both gardens, and included initial height and Block nested within Garden as fixed effects. The model also included source site locality as a random effect to account for regional differences; due to insufficient sample size of phenology data, we were unable to model differences among families.

We observed that families from warmer climates had earlier leaf emergence than those from cooler climates (Table S14). In addition, relative growth rates were higher in families with earlier leaf emergence across both gardens when controlling for climatic distance between source site and garden (Table S15). In addition, earlier leaf emergence date was associated with higher garden maximum temperatures during early spring when controlling for planting block within garden, year, and maternal seed source (February-April; *p* < 0.001, R^2^ marginal = 0.59, R^2^ conditional = 0.67). These findings indicate that length of growing season contributes to higher growth rates but is not the only mechanism that explains the difference between families of varying climatic origins.

As reported previously by Wright *et al.* (2021), progeny of trees from hotter environments had significantly earlier leaf emergence than trees from cooler environments. We found that earlier leaf emergence was correlated with higher growth rate and relative performance likely due to a longer growing season. Leaves also emerged earlier on average in years with warmer temperatures during the bud period (February-May), consistent with a previous study on valley oak (Gerst *et al.* 2017). In turn, we did not observe a tradeoff between a lengthened growing season and risk of frost damage; though late frosts occurred at the Placerville garden, we did not observe negative fitness effects of early leaf emergence. This finding suggests a possible future adaptational lag; that is, trees adapted to environments cooler than future conditions may continue to wait longer than necessary to put out leaves and in turn reduce their growth potential. However, the effects of frost damage on other aspects of plant health (e.g., leaf morphology, biomass and nutrient allocation) should be considered as well. Because we detected a pattern of local adaptation to mean winter temperature, other mechanisms likely impose a fitness penalty on families adapted to warmer environments than the garden. We thus find evidence that timing of leaf emergence partially predicts the fitness differences among valley oak trees from differing climates and may be a driver of future shifts toward lower fitness in cool-adapted populations.

#### **Appendix S9. Estimating baseline distribution of relative fitness for spatial predictions**

To construct the baseline relative fitness distribution rasters, we first estimated the current distribution of valley oak relative fitness across the entire species range using an additional “phenotype-environment association” GAM that related maternal trees’ 2024 relative fitness values to a series of environmental variables, fitted using the following formula:

**Eq. S4.** *RF_2024_ ~ α + s(T_max_) + s(T_min_) + s(Precip.) + s(Elevation) + te(Latitude, Longitude) + 𝜀*

where *α* is the intercept and 𝜀 is the error term. Terms within an s() parenthetical are smooth terms modeled using cubic regression splines, and terms within a te() parenthetical are tensor interaction splines, which model the additive effects of each term and their interaction. The environmental variables were summer maximum temperature, winter minimum temperature, summer precipitation, elevation above sea level, and the latitude × longitude interaction (to control for spatial autocorrelation). We used multiple environmental variables to provide a more realistic estimate of fitness values across the range than could be achieved with our models that only tested effects of one temperature variable. We then predicted relative fitness values from the phenotype-environment association across a raster stack of range-wide 1961-1990 climate data (AdaptWest 2022). We used 1961-1990 rasters for these analyses because 1961-1990 is the oldest period available in the AdaptWest dataset. We used AdaptWest rasters for both past and future environmental data to ensure that all rasters had the same spatial extent and resolution and that all environmental variables were estimated and interpolated the same way. We used the resulting raster as an interpolated distribution of relative fitness across the species range for predictive modeling. Spatial prediction was done using function predict(), which applies a given prediction function—in this case predict.gam from R package ‘mgcv’—across each pixel of a raster, in R package ‘terra’ version 1.8-54 (Hijmans *et al.* 2022).

#### **Appendix S10. Families differ primarily in growth, not survival**

Valley oak fitness differences were driven primarily by growth, not survival. Relative fitness models incorporating survival had lower explanatory power than our relative growth rate model, likely because most poorly performing trees remained small (<150 cm tall, with little secondary growth), rather than experiencing climate-induced mortality. This pattern has been observed in other oak provenance tests (Sáenz-Romero *et al.* 2017; Girard *et al.* 2022), which may be evidence of the resilience of this taxon. At the same time, we acknowledge that irrigation deliberately minimized mortality to retain sample sizes needed for genetic analyses. Despite this limitation, size is a robust indicator of high performance in trees and is strongly associated with lifetime fecundity (Wadgymar *et al.* 2024), including in oaks (Ramírez-Valiente *et al.* 2022). **While growth-based fitness differences could change as trees reach reproductive maturity** (St. Clair *et al.* 2020), the increasing divergence observed between five- and twelve-year-old trees makes it unlikely that early maladaptive trends will disappear later in life.

#### **Appendix S11. Effects of geography on fitness**

Data on maternal tree source site climates from the California Basin Characterization Model (Flint et al. 2013) were used to compare common garden growth outcomes against historical climatic conditions experienced by the maternal source trees. All measurements were 30-year averages of 1950-1981 data gridded at 270m pixel resolution. Temperature and precipitation data were used to calculate the BioClim variables (worldclim.org), and a principal components analysis was run on 10 climate variables (Maximum summer temperature, minimum winter temperature, maximum, minimum, and average annual temperatures, climatic water deficit, temperature seasonality, precipitation seasonality, precipitation of warmest quarter, precipitation of coldest quarter). The first PC axis (40.8% of variance explained) represented the spectrum between hotter, dryer sites and cooler, wetter sites, while the second PC axis (23.3% of variance explained) represented seasonal differences in temperature and precipitation (higher values indicate temperature-based seasonality, lower values indicate precipitation-based seasonality). For more detail on the climatic variables and PCAs, see Browne et al. 2019.

We tested the effects of geography on relative growth rate (RGR) and relative fitness. As we were interested in detecting geographic “hotspots” of overall successful families, we measured response of cumulative relative fitness and RGR (relative fitness in 2024; 2014-2024 cumulative RGR) to latitude, longitude, and the first two principal components describing variation in maternal seed source sites (higher PC1: hotter/dryer, higher PC2: temperature, not precipitation, drives seasonality; see “Maternal tree source climate data” section above). Gardens were analyzed separately. The growth rate models also included the fixed effects of initial height and garden block and the random effect of family. Explanatory variables were scaled as Z-scores. Phenology and geography models were fitted using the lmer() function in R package lme4 (Bates *et al.* 2015).

To test for the effect of geographic variation, including photoperiod, on cumulative relative fitness and RGR, we tested the relationship between 2024 relative fitness/2014-2024 RGR and longitude/latitude when controlling for the maternal climate PC vectors. We did not find significant relationships between relative fitness and latitude or longitude (no macroclimate effect), but the PC vectors, representing local climatic variation, significantly predicted relative fitness. Families from hotter, dryer, and more temperature-seasonal (as opposed to precipitation-seasonal) sites had significantly higher fitness (Placerville garden: *p*_PC1_ < 0.001, *p*_PC2_ = 0.01; Chico garden: *p*_PC1_ = 0.02, *p*_PC2_ < 0.001). Geographic variation was high even at the local climate scale, with trees from nearby sites showing varied fitness outcomes. Several maternal trees in the 95^th^ percentile for relative fitness were sourced from cooler sites than the gardens and many sites hotter than the gardens did not have high performing trees (Figure S6). Geographic effects on cumulative growth rates were slightly different. At the Placerville garden, latitude and PC1 both significantly predicted growth rate (*p*_latitude_ = 0.03, *p*_PC1_ < 0.001) as higher in families from hotter sites in the northern extent of the species range. At the Chico garden, only PC2 significantly predicted growth rate (*p*_PC2_ = 0.03), and families from more precipitation-seasonal sites had higher growth rate. Summaries are provided in Tables S15-S16. Finally, to visualize and interpret the geographic differences in relative fitness, we used two-dimensional kriging to interpolate relative fitness values between points within the hypothesized species range. Models were fitted using the gstat() function, and interpolated using the interpolate() function, in R package terra version 1.7-83 (Hijmans *et al.* 2022).

We did not find strong patterns of latitude or longitude predicting fitness such as reported for forest trees in Canada (e.g., Rehfeldt *et al.* 1999), nor did individuals clearly separate into distinct geographic units. Instead, we found high geographic variation in maternal tree fitness. Furthermore, though local climate was a significant predictor of relative fitness, several trees in the 95^th^ percentile for relative fitness were from sites up to 2 degrees cooler than the median garden temperature.

#### **Supplementary figures**

**Figure S1**: Correlation matrix of past summer maximum temperatures at source locations of 658 adult valley oak trees, in periods 1700-1800, 1900-2000, and 1951-1980. 1700-2000 data are from the CHELSA TraCE21k time series dataset (Karger *et al.* 2017; Karger *et al.* 2023); 1951-1980 data are from the California Basin Characterization Model (Flint *et al.* 2013).

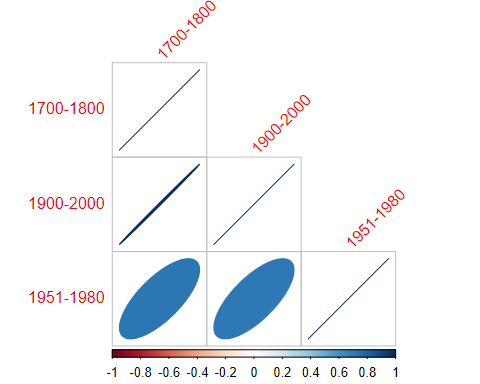

**Figure S2**: Distribution of maternal tree origin site and common garden temperatures (summer maximum; winter mean, °C), colored by elevation above sea level (m). Maternal origin sites are shown as dots; common gardens are shown as large triangles. Pearson correlation coefficient between summer maximum and winter mean temperature = -0.01. All origin sites with summer maximum temperature above 33°C have winter mean temperature between 7 and 10°C and there is a significant quadratic relationship between winter mean temperature and summer maximum temperature (F = 38.1, df = 656, p < 0.001). An elevational cline effect is apparent among sites with low summer and winter temperature (bottom right corner); low-elevation sites show wide variation in temperature. For maternal tree origin sites, temperature is the 1951-1980 average from the Basin Characterization Model (Flint *et al.* 2013). For common gardens, temperature data is the 2014-2024 average from PRISM (prism.oregonstate.edu).

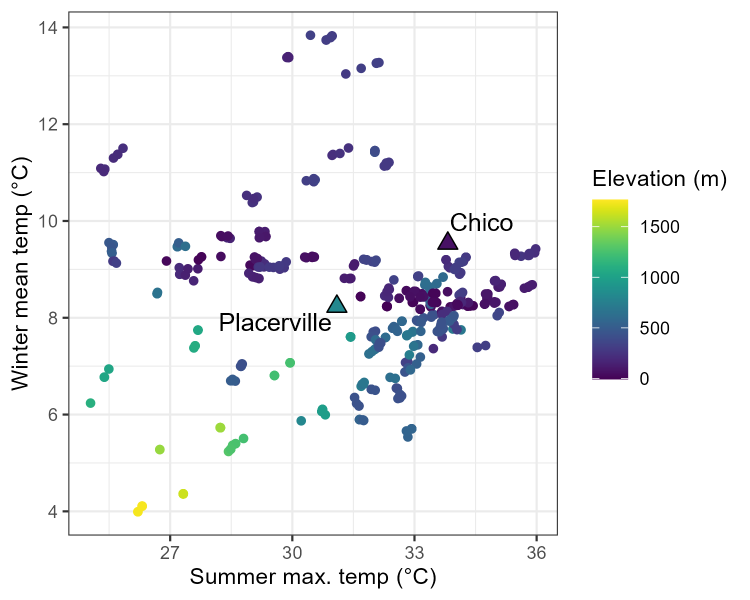

**Figure S3**: Total annual precipitation 2014-2024 in the Placerville, CA (blue) and Chico, CA (red) common gardens, with the mean of both gardens shown in the green dashed line. Data from PRISM (prism.oregonstate.edu). Refer to Methods for specific locations of the gardens.

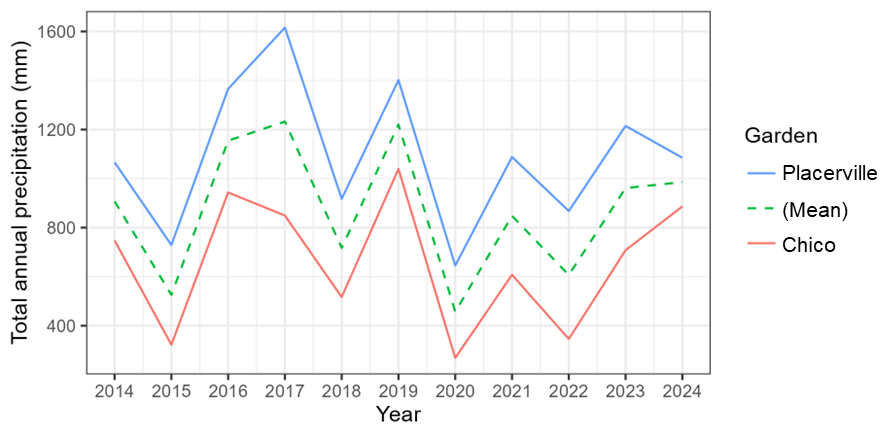

**Figure S4**: Ten-year cumulative trends in growth rate (left) and relative fitness (right) as a function of temperature transfer distance (maximum monthly temperature averaged across June-August) between maternal tree source location and common garden. Solid curves denote generalized additive models fit to empirical data, and dashed curves denote extrapolation of the models using Gaussian process regression. Shaded regions represent 95% confidence intervals. The peak in each extrapolated curve represents the theoretical species-wide trait optimum. Vertical dashed line in each plot denotes where source temperature equals garden temperature; vertical solid lines denote estimated average temperature during the Last Glacial Maximum (~21 kya; Karger *et al.* 2023) and predicted average temperature in year 2100 under a stabilization climate scenario (RCP 4.5; Intergovernmental Panel on Climate 2023). The area to the left of the zero line contains individuals from warmer historical climates than the garden, while the area to the right contains individuals from cooler historical climates.

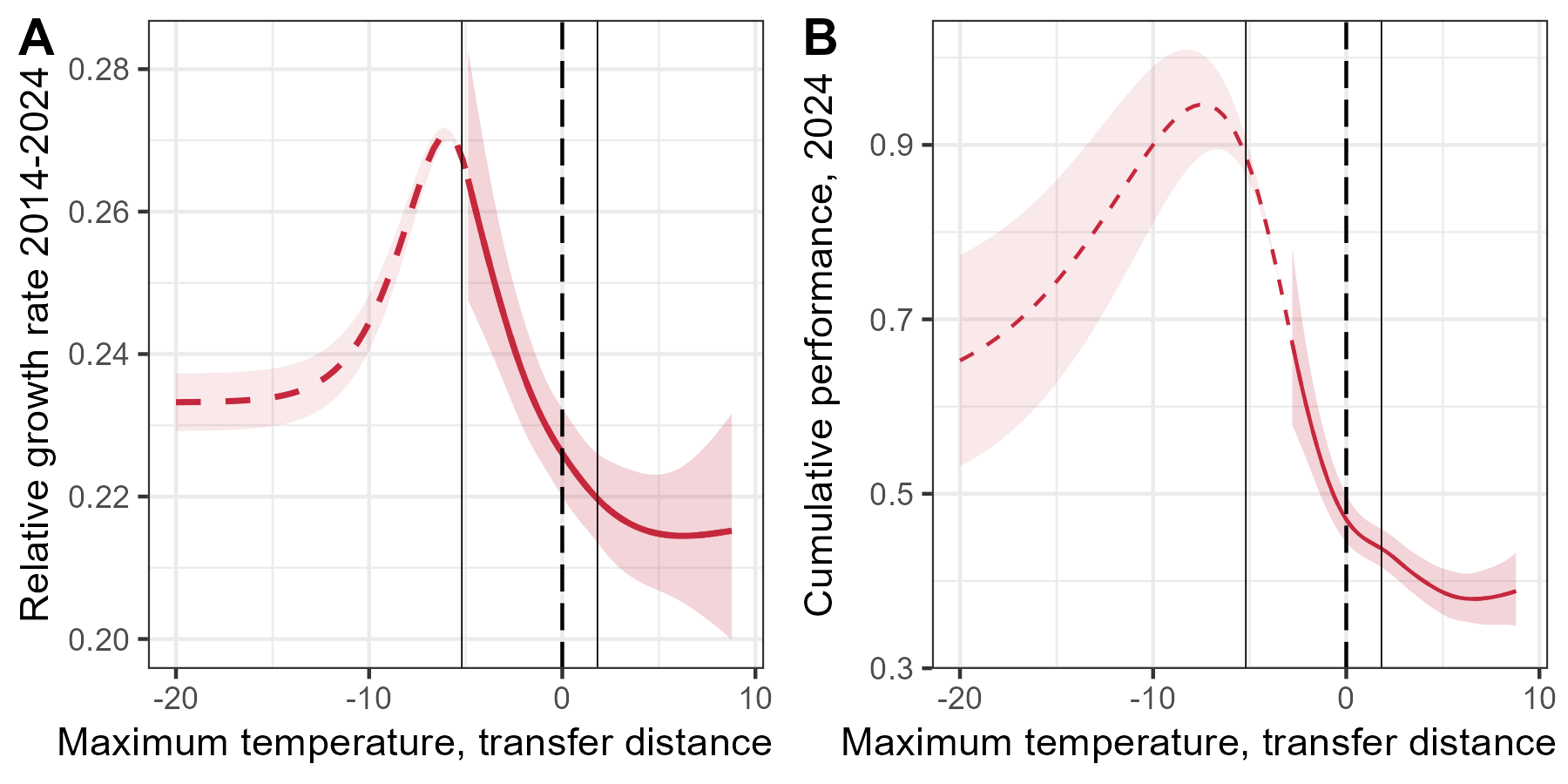

**Figure S5**: Yearly trends in valley oak growth rate as a function of transfer distance between maternal location and common garden, arranged and color-coded in order of cooler to warmer temperatures. Solid colored curves represent empirically derived generalized additive model predictions of single-year relative growth rate as a function of yearly temperature transfer distance (maximum monthly temperature averaged across June-August) across the period of tree growth in the common gardens, 2015-2024 (2018 is excluded due to incomplete data). Long-dashed curves represent extrapolations of the model using Gaussian process regression. Thin dotted curves represent non-significant relationships; thicker dotted curves represent marginally significant relationships (p < 0.1). Vertical dashed line in each plot denotes where source temperature equals garden temperature; vertical solid lines denote estimated average temperature during the Last Glacial Maximum (~21 kya; Karger *et al.* 2023) and predicted average temperature in year 2100 under a stabilization climate scenario (RCP 4.5; Intergovernmental Panel on Climate 2023). The area to the left of the zero line contains individuals from warmer historical climates than the garden, while the area to the right contains individuals from cooler historical climates.

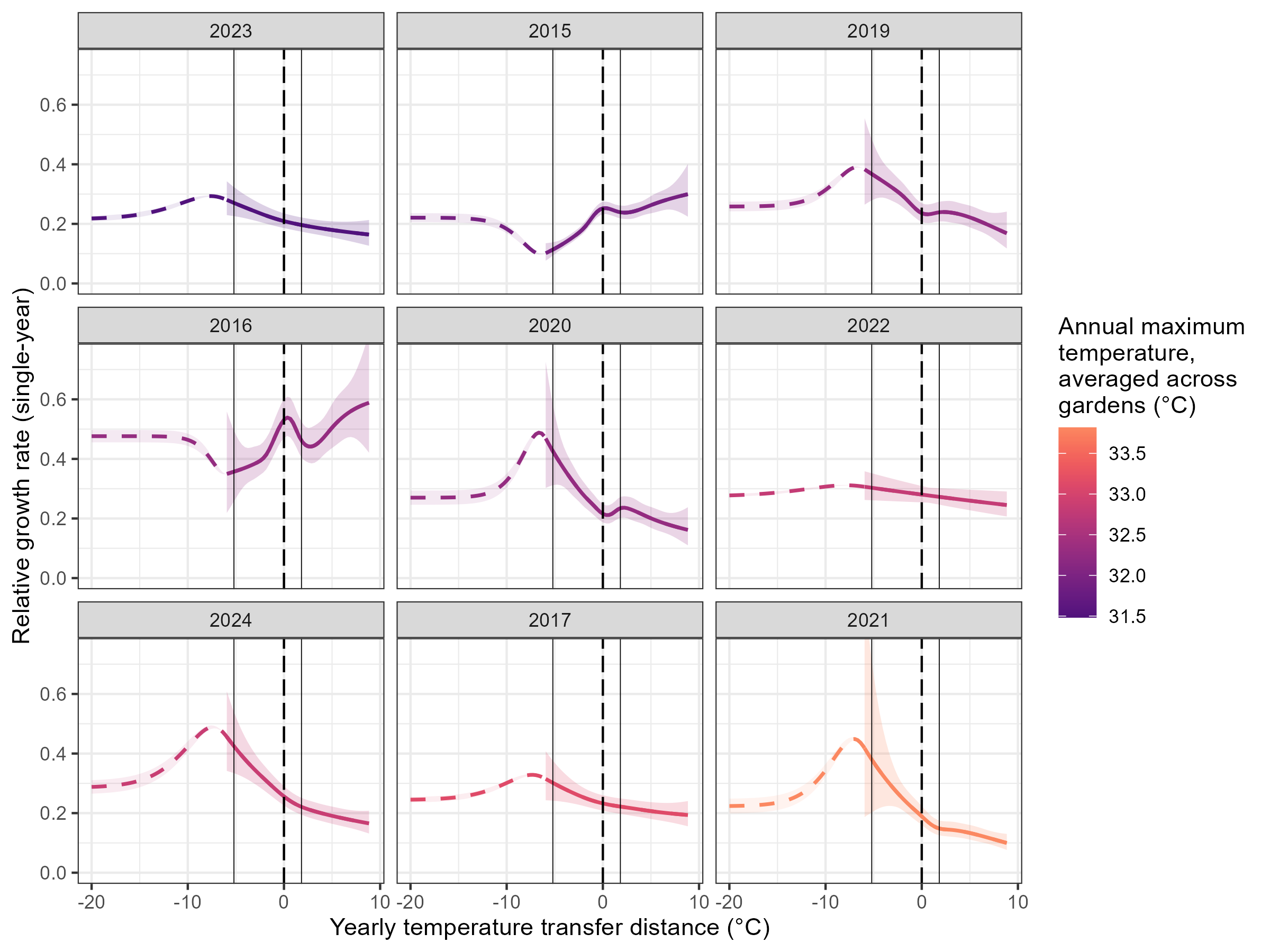

**Figure S6**: Geographic distribution of relative fitness of maternal seed sources measured in the common gardens (left: Placerville garden, right: Chico garden) with point locations indicated by blue or red dot surrounded by 2500-meter interpolated buffer zones. Color gradient of buffer zone shows family mean relative fitness in the common gardens in 2024.

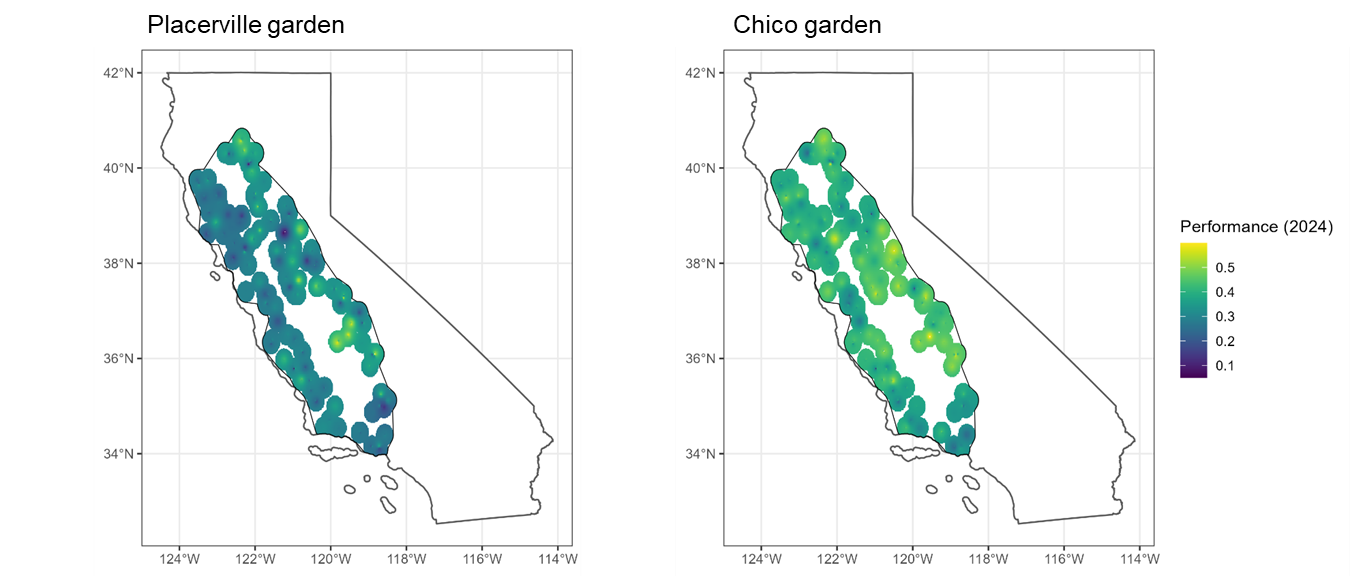

#### **Supplementary tables**

**Table S1**: Summary of growth variables measured on valley oak progeny in the common gardens in each year. Starting in 2018, some individuals were too tall to efficiently measure to their full heights so cutoff thresholds (shown in parentheses) were imposed.

| **Year** | **Growth variables measured** |
| --- | --- |
| 2013 | Height, basal diameter |
| 2014 | Height, basal diameter |
| 2015 | Height, basal diameter |
| 2016 | Height |
| 2017 | Height |
| 2018 | Height (<4 m), DBH (haphazard subset), basal diameter (haphazard subset) |
| 2019 | Height (<4 m), DBH (all trees above 150 cm height) |
| 2020 | DBH or basal diameter (IFG: all; CSO: 4/5 blocks) |
| 2021 | Height (<3 m), DBH (all trees above 150 cm height); basal diameter (removed trees), actual height (removed trees) |
| 2022 | Height (<2 m), DBH (all trees above 150 cm height) |
| 2023 | Height (<2.5 m IFG; <2 m CSO), DBH (all trees above 150 cm height), actual height (haphazard subset) |
| 2024 | Height (all IFG; <2m CSO), DBH (all trees above 150 cm height) |

**Table S2:** Tree height is genetically differentiated by maternal family. Two-way ANOVA by family and garden. Height significantly differs among families and gardens, but the family*garden interaction term is not significant, i.e., the same genotypes tend to grow better or worse in each garden. The significant effect of planting block is included to account for known environmental differences within the gardens.

|  | *df* | *F* | *p* |
| --- | --- | --- | --- |
| Garden | **1** | **467.73** | **< 0.001** |
| Family | **630** | **1.80** | **< 0.001** |
| Block | **4** | **63.25** | **< 0.001** |
| Garden:Family | 620 | 1.05 | 0.24 |
| Residuals | 2417 |  |  |

**Table S3:** Tree relative growth rate is genetically differentiated by maternal family across years. Two-way repeated measures ANOVA tests on yearly relative growth rate by family. Between-groups comparisons test whether individual trees differ from one another, while within-groups comparisons test whether patterns within the same individual trees change over time. The planting block term is included to account for known environmental differences within gardens. **A:** Placerville garden; **B:** Chico garden.

|  |  | A. Placerville garden | | | B. Chico garden | | | |
| --- | --- | --- | --- | --- | --- | --- | --- | --- |
|  |  | *df* | *t* | *p* | | *df* | *t* | *p* |
| Between-groups | Family | **621** | **2.8** | **< 0.001** | | **624** | **2.0** | **< 0.001** |
|  | Year | **9** | **18.1** | **0.006** | | **9** | **27.2** | **< 0.001** |
|  | Planting block | **4** | **96.4** | **< 0.001** | | **4** | **89.1** | **< 0.001** |
|  | Family:Year | **982** | **2.5** | **< 0.001** | | 1,038 | 1.3 | 0.2 |
|  | Residuals | 39 |  |  | | 42 |  |  |
| Within-groups | Year | **9** | **473.3** | **< 0.001** | | **9** | **1,364.7** | **< 0.001** |
|  | Family:Year | 5,329 | 1.0 | 0.6 | | 5,354 | 0.8 | 1.0 |
|  | Residuals | 5,900 |  |  | | 6,345 |  |  |

**Table S4**: Summary of GAM estimating cumulative effect of summer maximum temperature transfer distance on relative growth rate, 2014-2024. Root mean square error from 10-fold model cross-validation is reported below. The planting block term (nested within site) is included to account for known environmental differences within gardens.

| **Component** | **Term** | **Est.** | **Std Error** | **t** | **p** |  |
| --- | --- | --- | --- | --- | --- | --- |
| A. parametric coefficients | (Intercept) | -1.44 | 0.01 | -112.66 | < 0.001 | *** |
|  | SiteBlockChico2 | -0.07 | 0.02 | -3.50 | < 0.001 | *** |
|  | SiteBlockChico3 | -0.13 | 0.02 | -7.83 | < 0.001 | *** |
|  | SiteBlockChico4 | -0.16 | 0.02 | -9.59 | < 0.001 | *** |
|  | SiteBlockChico5 | -0.11 | 0.02 | -6.48 | < 0.001 | *** |
|  | SiteBlockIFG1 | -0.26 | 0.02 | -13.66 | < 0.001 | *** |
|  | SiteBlockIFG2 | -0.29 | 0.02 | -15.70 | < 0.001 | *** |
|  | SiteBlockIFG3 | -0.32 | 0.02 | -16.67 | < 0.001 | *** |
|  | SiteBlockIFG4 | -0.30 | 0.02 | -15.95 | < 0.001 | *** |
|  | SiteBlockIFG5 | -0.39 | 0.02 | -19.48 | < 0.001 | *** |
| **Component** | **Term** |  |  |  |  |  |
| B. smooth terms | Smooth term (2014 height) | 5.42 | 6.74 | 392.04 | < 0.001 | *** |
|  | Smooth term (Tdiff, summer max) | 3.00 | 3.45 | 14.32 | < 0.001 | *** |
|  | Smooth term (Locality) | 53.23 | 94.00 | 1.48 | < 0.001 | *** |
|  | Smooth term (Family) | 23.50 | 625.00 | 0.04 | 0.25 |  |
| Signif. codes: 0 <= '***' < 0.001 < '**' < 0.01 < '*' < 0.05 | | | | | | |
| Adjusted R-squared: 0.603, Deviance explained 0.553 | | | | | | |
| fREML : -5262.909, Scale est: 0.0113, N: 3351 | | | | | | |

10-fold cross-validation RMSE: 1.86 ± 0.02

**Table S5**: Summary of GAM estimating cumulative effect of summer maximum temperature transfer distance on relative fitness, 2014-2024. Root mean square error from 10-fold model cross-validation is reported below.

| **Component** | **Term** | **Est.** | **Std Error** | **t** | **p** |  |
| --- | --- | --- | --- | --- | --- | --- |
| A. parametric coefficients | (Intercept) | -0.85 | 0.02 | -45.71 | < 0.001 | *** |
|  | Garden: IFG | -0.35 | 0.03 | -13.27 | < 0.001 | *** |
| **Component** | **Term** |  |  |  |  |  |
| B. smooth terms | Smooth term (Tdiff, summer max) | 3.57 | 3.86 | 15.69 | < 0.001 | *** |
|  | Smooth term (2014 relative fitness) | 3.14 | 3.92 | 8.55 | < 0.001 | *** |
|  | Smooth term (Family) | < 0.001 | 621.00 | < 0.001 | 0.57 |  |
|  | Smooth term (Locality) | 43.16 | 94.00 | 0.92 | < 0.001 | *** |
| Signif. codes: 0 <= '***' < 0.001 < '**' < 0.01 < '*' < 0.05 | | | | | | |
| Adjusted R-squared: 0.236, Deviance explained 0.265 | | | | | | |
| fREML : -742.512, Scale est: 0.0493, N: 1238 | | | | | | |

10-fold cross-validation RMSE: 1.40 ± 0.02

**Table S6**: Summary of GAM estimating cumulative effect of averaged summer maximum temperature transfer distance on relative growth rate, 2014-2024. Root mean square error from 10-fold model cross-validation is reported below. The planting block term (nested within site) is included to account for known environmental differences within gardens.

| **Component** | **Term** | **Est.** | **Std Error** | **t** | **p** |  |
| --- | --- | --- | --- | --- | --- | --- |
| A. parametric coefficients | (Intercept) | -1.44 | 0.01 | -110.95 | < 0.001 | *** |
|  | SiteBlockChico2 | -0.07 | 0.02 | -3.49 | < 0.001 | *** |
|  | SiteBlockChico3 | -0.13 | 0.02 | -7.84 | < 0.001 | *** |
|  | SiteBlockChico4 | -0.16 | 0.02 | -9.59 | < 0.001 | *** |
|  | SiteBlockChico5 | -0.11 | 0.02 | -6.49 | < 0.001 | *** |
|  | SiteBlockIFG1 | -0.26 | 0.02 | -13.73 | < 0.001 | *** |
|  | SiteBlockIFG2 | -0.30 | 0.02 | -15.73 | < 0.001 | *** |
|  | SiteBlockIFG3 | -0.32 | 0.02 | -16.67 | < 0.001 | *** |
|  | SiteBlockIFG4 | -0.31 | 0.02 | -15.97 | < 0.001 | *** |
|  | SiteBlockIFG5 | -0.40 | 0.02 | -19.45 | < 0.001 | *** |
| **Component** | **Term** |  |  |  |  |  |
| B. smooth terms | Smooth term (2014 height) | 5.43 | 6.77 | 390.71 | < 0.001 | *** |
|  | Smooth term (Tdiff, summer averaged max) | 2.97 | 3.44 | 14.19 | < 0.001 | *** |
|  | Smooth term (Locality) | 53.55 | 94.00 | 1.49 | < 0.001 | *** |
|  | Smooth term (Family) | 23.49 | 625.00 | 0.04 | 0.25 |  |
| Signif. codes: 0 <= '***' < 0.001 < '**' < 0.01 < '*' < 0.05 | | | | | | |
| Adjusted R-squared: 0.603, Deviance explained 0.553 | | | | | | |
| fREML : -5262.774, Scale est: 0.0113, N: 3351 | | | | | | |
| 10-fold cross-validation RMSE: 1.86 ± 0.01 | | | | | | |

**Table S7**: Summary of GAM estimating cumulative effect of averaged summer maximum temperature transfer distance on relative fitness, 2014-2024. Root mean square error from 10-fold model cross-validation is reported below.

| **Component** | **Term** | **Est.** | **Std Error** | **t** | **p** |  |
| --- | --- | --- | --- | --- | --- | --- |
| A. parametric coefficients | (Intercept) | -0.84 | 0.02 | -43.88 | < 0.001 | *** |
|  | Garden: IFG | -0.37 | 0.03 | -13.16 | < 0.001 | *** |
| **Component** | **Term** |  |  |  |  |  |
| B. smooth terms | Smooth term (Tdiff, summer averaged max) | 4.22 | 5.10 | 11.91 | < 0.001 | *** |
|  | Smooth term (2014 relative fitness) | 3.15 | 3.93 | 8.49 | < 0.001 | *** |
|  | Smooth term (Family) | < 0.001 | 621.00 | < 0.001 | 0.58 |  |
|  | Smooth term (Locality) | 43.29 | 94.00 | 0.92 | < 0.001 | *** |
| Signif. codes: 0 <= '***' < 0.001 < '**' < 0.01 < '*' < 0.05 | | | | | | |
| Adjusted R-squared: 0.235, Deviance explained 0.265 | | | | | | |
| fREML : -742.213, Scale est: 0.0493, N: 1238 | | | | | | |

10-fold cross-validation RMSE: 1.40 ± 0.02

**Table S8**: Summary of generalized additive model estimating cumulative effect of mean winter temperature transfer distance on relative growth rate, 2014-2024. Root mean square error from 10-fold model cross-validation is reported below. The planting block term (nested within site) is included to account for known environmental differences within gardens.

| **Component** | **Term** | **Est.** | **Std Error** | **t** | **p** |  |
| --- | --- | --- | --- | --- | --- | --- |
| A. parametric coefficients | (Intercept) | -1.46 | 0.01 | -111.83 | < 0.001 | *** |
|  | SiteBlockChico2 | -0.07 | 0.02 | -3.48 | < 0.001 | *** |
|  | SiteBlockChico3 | -0.13 | 0.02 | -7.69 | < 0.001 | *** |
|  | SiteBlockChico4 | -0.16 | 0.02 | -9.57 | < 0.001 | *** |
|  | SiteBlockChico5 | -0.11 | 0.02 | -6.41 | < 0.001 | *** |
|  | SiteBlockIFG1 | -0.22 | 0.02 | -11.48 | < 0.001 | *** |
|  | SiteBlockIFG2 | -0.25 | 0.02 | -13.46 | < 0.001 | *** |
|  | SiteBlockIFG3 | -0.28 | 0.02 | -14.42 | < 0.001 | *** |
|  | SiteBlockIFG4 | -0.26 | 0.02 | -13.81 | < 0.001 | *** |
|  | SiteBlockIFG5 | -0.35 | 0.02 | -17.45 | < 0.001 | *** |
| **Component** | **Term** |  |  |  |  |  |
| B. smooth terms | Smooth term (2014 height) | 5.24 | 6.52 | 400.27 | < 0.001 | *** |
|  | Smooth term (Tdiff, winter mean) | 3.05 | 3.44 | 7.16 | < 0.001 | *** |
|  | Smooth term (Locality) | 58.20 | 94.00 | 1.88 | < 0.001 | *** |
|  | Smooth term (Family) | 18.60 | 625.00 | 0.03 | 0.29 |  |
| Signif. codes: 0 <= '***' < 0.001 < '**' < 0.01 < '*' < 0.05 | | | | | | |
| Adjusted R-squared: 0.602, Deviance explained 0.552 | | | | | | |
| fREML : -5254.382, Scale est: 0.0114, N: 3351 | | | | | | |

10-fold cross-validation RMSE: 1.86 ± 0.01

**Table S9**: Summary of GAM estimating cumulative effect of mean winter temperature transfer distance on relative fitness, 2014-2024. Root mean square error from 10-fold model cross-validation is reported below.

| **Component** | **Term** | **Est.** | **Std Error** | **t** | **p** |  |
| --- | --- | --- | --- | --- | --- | --- |
| A. parametric coefficients | (Intercept) | -0.89 | 0.02 | -47.12 | < 0.001 | *** |
|  | Garden: IFG | -0.25 | 0.02 | -11.80 | < 0.001 | *** |
| **Component** | **Term** |  |  |  |  |  |
| B. smooth terms | Smooth term (Tdiff, winter mean) | 2.86 | 3.25 | 3.58 | 0.01 | * |
|  | Smooth term (2014 relative fitness) | 2.95 | 3.69 | 9.01 | < 0.001 | *** |
|  | Smooth term (Family) | < 0.001 | 621.00 | < 0.001 | 0.68 |  |
|  | Smooth term (Locality) | 53.39 | 94.00 | 1.46 | < 0.001 | *** |
| Signif. codes: 0 <= '***' < 0.001 < '**' < 0.01 < '*' < 0.05 | | | | | | |
| Adjusted R-squared: 0.222, Deviance explained 0.260 | | | | | | |
| fREML : -724.015, Scale est: 0.0503, N: 1238 | | | | | | |

10-fold cross-validation RMSE: 1.40 ± 0.02

**Table S10**: Summary of GAM measuring interannual variation in effects of annual maximum temperature transfer distance on individual relative growth rates over time. Root mean square error from 10-fold model cross-validation is reported below. The planting block term (nested within site) is included to account for known environmental differences within gardens. Terms beginning with “s(tdiffyr)” represent the slope of the temperature-fitness relationship in each study year.

| **Component** | **Term** | **Est.** | **Std Error** | **t** | **p** |  |
| --- | --- | --- | --- | --- | --- | --- |
| A. parametric coefficients | (Intercept) | -1.32 | 0.02 | -61.18 | < 0.001 | *** |
|  | Year2016 | 0.69 | 0.02 | 36.66 | < 0.001 | *** |
|  | Year2017 | -0.02 | 0.02 | -0.65 | 0.52 |  |
|  | Year2019 | 0.05 | 0.03 | 2.01 | 0.04 | * |
|  | Year2020 | 0.03 | 0.03 | 1.05 | 0.29 |  |
|  | Year2021 | -0.27 | 0.03 | -8.74 | < 0.001 | *** |
|  | Year2022 | 0.17 | 0.03 | 6.38 | < 0.001 | *** |
|  | Year2023 | -0.07 | 0.03 | -2.60 | 0.01 | ** |
|  | Year2024 | 0.21 | 0.03 | 6.12 | < 0.001 | *** |
|  | SiteBlockChico2 | -0.17 | 0.02 | -7.22 | < 0.001 | *** |
|  | SiteBlockChico3 | -0.26 | 0.02 | -12.78 | < 0.001 | *** |
|  | SiteBlockChico4 | -0.33 | 0.02 | -16.53 | < 0.001 | *** |
|  | SiteBlockChico5 | -0.27 | 0.02 | -14.11 | < 0.001 | *** |
|  | SiteBlockIFG1 | -0.40 | 0.02 | -17.91 | < 0.001 | *** |
|  | SiteBlockIFG2 | -0.42 | 0.02 | -18.85 | < 0.001 | *** |
|  | SiteBlockIFG3 | -0.46 | 0.02 | -19.42 | < 0.001 | *** |
|  | SiteBlockIFG4 | -0.41 | 0.02 | -18.42 | < 0.001 | *** |
|  | SiteBlockIFG5 | -0.49 | 0.02 | -20.97 | < 0.001 | *** |
| **Component** | **Term** |  |  |  |  |  |
| B. smooth terms | s(tdiffyr):Year2015 | 5.47 | 6.45 | 38.67 | < 0.001 | *** |
|  | s(tdiffyr):Year2016 | 6.88 | 7.84 | 14.13 | < 0.001 | *** |
|  | s(tdiffyr):Year2017 | 2.64 | 3.30 | 8.05 | < 0.001 | *** |
|  | s(tdiffyr):Year2019 | 4.20 | 5.16 | 13.49 | < 0.001 | *** |
|  | s(tdiffyr):Year2020 | 6.74 | 7.77 | 12.73 | < 0.001 | *** |
|  | s(tdiffyr):Year2021 | 4.51 | 5.38 | 29.52 | < 0.001 | *** |
|  | s(tdiffyr):Year2022 | 1.00 | 1.00 | 5.81 | 0.02 | * |
|  | s(tdiffyr):Year2023 | 1.97 | 2.47 | 13.44 | < 0.001 | *** |
|  | s(tdiffyr):Year2024 | 3.38 | 4.07 | 40.85 | < 0.001 | *** |
|  | Smooth term (Year-1 height) | 12.58 | 13.56 | 512.39 | < 0.001 | *** |
|  | Smooth term (2014 height) | 3.17 | 3.96 | 5.09 | < 0.001 | *** |
|  | Smooth term (Family) | 167.58 | 626.00 | 0.39 | < 0.001 | *** |
|  | Smooth term (Locality) | 48.07 | 94.00 | 1.63 | < 0.001 | *** |
| Signif. codes: 0 <= '***' < 0.001 < '**' < 0.01 < '*' < 0.05 | | | | | | |
| Adjusted R-squared: 0.601, Deviance explained 0.580 | | | | | | |
| fREML : -15971.051, Scale est: 0.320, N: 24156 | | | | | | |

10-fold cross-validation RMSE: 1.92 ± 0.01

**Table S11**: Summaries of ANCOVAs on the interactions between maternal origin maximum temperature (T_origin_) and annual maximum garden temperature (T_garden_) of the current and previous study year (T_garden;yr-1_). Planting block is included in the models to account for known environmental differences within gardens. A: Placerville garden; B: Chico garden.

|  | | A. Placerville garden | | | | | | B. Chico garden | | | | |
| --- | --- | --- | --- | --- | --- | --- | --- | --- | --- | --- | --- | --- |
|  | *df* | | *SS* | *MS* | *F* | *p* | *df* | | *SS* | *MS* | *F* | *p* |
| T_origin_ | **1** | | **0.6** | **0.6** | **20.2** | **< 0.001** | **1** | | **0.3** | **0.3** | **22.4** | **< 0.001** |
| T_garden_ | **1** | | **9.1** | **9.1** | **311.2** | **< 0.001** | **1** | | **0.4** | **0.4** | **26.4** | **< 0.001** |
| T_garden;yr-1_ | **1** | | **9.2** | **9.2** | **313.2** | **< 0.001** | **1** | | **1.6** | **1.6** | **104.7** | **< 0.001** |
| T_origin_ × T_garden_ | 1 | | 0.1 | 0.1 | 3.5 | 0.06 | **1** | | **0.1** | **0.1** | **7.5** | **0.006** |
| T_origin_ × T_garden;yr-1_ | **1** | | **0.1** | **0.1** | **4.7** | **0.03** | 1 | | <0.05 | <0.05 | 2.3 | 0.13 |
| Planting block | **4** | | **0.9** | **0.2** | **7.7** | **< 0.001** | **4** | | **2.1** | **0.5** | **34.3** | **< 0.001** |
| Residuals | 10,145 | | 296.9 | <0.05 |  |  | 8,048 | | 124.1 | <0.05 |  |  |

**Table S12**: Summary of GAM measuring interannual variation in effects of averaged summer maximum temperature transfer distance on individual relative growth rates over time. Root mean square error from 10-fold model cross-validation is reported below. The planting block term (nested within site) is included to account for known environmental differences within gardens. The planting block term (nested within site) is included to account for known environmental differences within gardens. Terms beginning with “s(tdiffyr2)” represent the slope of the temperature-fitness relationship in each study year.

| **Component** | **Term** | **Est.** | **Std Error** | **t** | **p** |  |
| --- | --- | --- | --- | --- | --- | --- |
| A. parametric coefficients | (Intercept) | -1.38 | 0.02 | -67.17 | < 0.001 | *** |
|  | Year2016 | 0.74 | 0.02 | 43.69 | < 0.001 | *** |
|  | Year2017 | 0.06 | 0.02 | 2.74 | 0.01 | ** |
|  | Year2019 | 0.14 | 0.02 | 6.13 | < 0.001 | *** |
|  | Year2020 | 0.10 | 0.02 | 4.06 | < 0.001 | *** |
|  | Year2021 | -0.15 | 0.03 | -4.61 | < 0.001 | *** |
|  | Year2022 | 0.25 | 0.03 | 10.02 | < 0.001 | *** |
|  | Year2023 | -0.05 | 0.03 | -1.67 | 0.10 | . |
|  | Year2024 | 0.15 | 0.03 | 5.66 | < 0.001 | *** |
|  | SiteBlockChico2 | -0.17 | 0.02 | -7.30 | < 0.001 | *** |
|  | SiteBlockChico3 | -0.26 | 0.02 | -12.78 | < 0.001 | *** |
|  | SiteBlockChico4 | -0.33 | 0.02 | -16.47 | < 0.001 | *** |
|  | SiteBlockChico5 | -0.27 | 0.02 | -13.96 | < 0.001 | *** |
|  | SiteBlockIFG1 | -0.43 | 0.02 | -19.21 | < 0.001 | *** |
|  | SiteBlockIFG2 | -0.46 | 0.02 | -20.07 | < 0.001 | *** |
|  | SiteBlockIFG3 | -0.50 | 0.02 | -20.63 | < 0.001 | *** |
|  | SiteBlockIFG4 | -0.45 | 0.02 | -19.76 | < 0.001 | *** |
|  | SiteBlockIFG5 | -0.53 | 0.02 | -22.22 | < 0.001 | *** |
| **Component** | **Term** |  |  |  |  |  |
| B. smooth terms | s(tdiffyr2):Year2015 | 5.74 | 6.75 | 31.93 | < 0.001 | *** |
|  | s(tdiffyr2):Year2016 | 6.52 | 7.51 | 11.68 | < 0.001 | *** |
|  | s(tdiffyr2):Year2017 | 2.74 | 3.42 | 11.98 | < 0.001 | *** |
|  | s(tdiffyr2):Year2019 | 5.52 | 6.55 | 17.53 | < 0.001 | *** |
|  | s(tdiffyr2):Year2020 | 6.28 | 7.29 | 21.00 | < 0.001 | *** |
|  | s(tdiffyr2):Year2021 | 4.51 | 5.34 | 34.84 | < 0.001 | *** |
|  | s(tdiffyr2):Year2022 | 1.00 | 1.00 | 15.20 | < 0.001 | *** |
|  | s(tdiffyr2):Year2023 | 2.31 | 2.89 | 23.01 | < 0.001 | *** |
|  | s(tdiffyr2):Year2024 | 3.18 | 3.93 | 54.52 | < 0.001 | *** |
|  | Smooth term (Year-1 height) | 12.58 | 13.56 | 515.15 | < 0.001 | *** |
|  | Smooth term (2014 height) | 3.11 | 3.88 | 4.93 | < 0.001 | *** |
|  | Smooth term (Family) | 166.82 | 626.00 | 0.38 | < 0.001 | *** |
| Signif. codes: 0 <= '***' < 0.001 < '**' < 0.01 < '*' < 0.05 | | | | | | |
| Adjusted R-squared: 0.606, Deviance explained 0.582 | | | | | | |
| fREML : -16028.427, Scale est: 0.318, N: 24156 | | | | | | |

10-fold cross-validation RMSE: 1.92 ± 0.01

**Table S13**: Date of first budburst of common garden valley oak trees differs among years and by family nested within locality (random effects). Estimates are estimated budburst date in each year (relative to 2015 budburst date estimate). σ2 = within-group variance; τ00 = between-group variance; ICC = intra-class correlation coefficient.

|  | **Date of first budburst** | | |
| --- | --- | --- | --- |
| *Predictors* | *Estimates* | *CI* | *p* |
| (Intercept) | -11965.64 | -12195.63 – -11735.64 | **<0.001** |
| SiteBlock [Chico2] | 1.66 | 0.55 – 2.77 | **0.003** |
| SiteBlock [Chico3] | 5.32 | 4.42 – 6.21 | **<0.001** |
| SiteBlock [Chico4] | 1.27 | 0.37 – 2.16 | **0.006** |
| SiteBlock [Chico5] | 1.42 | 0.46 – 2.38 | **0.004** |
| SiteBlock [IFG1] | 5.48 | 4.57 – 6.39 | **<0.001** |
| SiteBlock [IFG2] | 6.09 | 5.15 – 7.03 | **<0.001** |
| SiteBlock [IFG3] | 7.01 | 6.04 – 7.98 | **<0.001** |
| SiteBlock [IFG4] | 6.45 | 5.51 – 7.38 | **<0.001** |
| SiteBlock [IFG5] | 6.69 | 5.68 – 7.70 | **<0.001** |
| Year | 5.97 | 5.86 – 6.09 | **<0.001** |
| **Random Effects** | | | |
| σ^2^ | 82.53 | | |
| τ_00_ _Family:Locality_ | 3.93 | | |
| τ_00_ _Locality_ | 12.61 | | |
| ICC | 0.17 | | |
| N _Family_ | 627 | | |
| N _Locality_ | 95 | | |
| Observations | 8839 | | |
| Marginal R^2^ / Conditional R^2^ | 0.530 / 0.608 | | |

**Table S14**: Linear mixed model of valley oak trees’ individual relative growth rates as a function of year, temperature transfer distance, source site locality, and phenology. Estimates are relative growth rates estimated for each model term. σ2 = within-group variance; τ00 = between-group variance; ICC = intra-class correlation coefficient.

|  | **Relative growth rate, single-year** | | |
| --- | --- | --- | --- |
| *Predictors* | *Estimates* | *CI* | *p* |
| (Intercept) | 231.13 | 216.52 – 245.73 | **<0.001** |
| Date of first budburst | 0.01 | -0.00 – 0.02 | 0.071 |
| Year | -0.11 | -0.12 – -0.11 | **<0.001** |
| Temperature transfer distance | 0.00 | -0.01 – 0.01 | 0.622 |
| SiteBlockChico2 | -0.28 | -0.33 – -0.23 | **<0.001** |
| SiteBlockChico3 | -0.07 | -0.11 – -0.03 | **<0.001** |
| SiteBlockChico4 | -0.21 | -0.25 – -0.18 | **<0.001** |
| SiteBlockChico5 | -0.40 | -0.44 – -0.35 | **<0.001** |
| SiteBlockIFG1 | -0.39 | -0.43 – -0.34 | **<0.001** |
| SiteBlockIFG2 | -0.35 | -0.39 – -0.31 | **<0.001** |
| SiteBlockIFG3 | -0.29 | -0.34 – -0.25 | **<0.001** |
| SiteBlockIFG4 | -0.38 | -0.42 – -0.34 | **<0.001** |
| SiteBlockIFG5 | -0.40 | -0.45 – -0.36 | **<0.001** |
| **Random Effects** | | | |
| σ^2^ | 0.16 | | |
| τ_00_ _Locality_ | 0.00 | | |
| ICC | 0.00 | | |
| N _Locality_ | 95 | | |
| Observations | 8839 | | |
| Marginal R^2^ / Conditional R^2^ | 0.257 / 0.259 | | |

**Table S15:** Summary of phenotype-environment association fitted by generalized additive model, used to estimate baseline distribution of relative fitness for spatial prediction (Appendix S10). This model includes additional environmental variables relative to the previous GAMs (winter mean temperature and annual precipitation) to provide a more detailed prediction of geographic variation in relative fitness. In addition, latitude, longitude, and the latitude*longitude interaction are included to account for spatial autocorrelation.

| Component | Term | Est. | Std Error | t | p |  |
| --- | --- | --- | --- | --- | --- | --- |
| A. parametric coefficients | (Intercept) | 0.37 | 0.02 | 16.43 | < 0.001 | *** |
| Component | Term |  |  |  |  |  |
| B. smooth terms | Smooth term (Summer max. temp., 1961-1990 mean) | 2.19 | 2.74 | 1.71 | 0.24 |  |
|  | Smooth term (Winter mean temp., 1961-1990 mean) | 1.00 | 1.00 | 0.85 | 0.36 |  |
|  | Smooth term (Summer precip., 1961-1990 mean) | 1.00 | 1.00 | 0.06 | 0.81 |  |
|  | Smooth term (Elevation above sea level) | 1.00 | 1.00 | 26.76 | < 0.001 | *** |
|  | Smooth term (Latitude) | 4.33 | 5.37 | 7.98 | < 0.001 | *** |
|  | Smooth term (Longitude) | 3.48 | 4.42 | 8.26 | < 0.001 | *** |
|  | Tensor interaction (Latitude:Longitude) | 1.00 | 1.00 | 0.65 | 0.42 |  |
| Signif. codes: 0 <= '***' < 0.001 < '**' < 0.01 < '*' < 0.05 | | | | | | |
| Adjusted R-squared: 0.0757, Deviance explained 0.0859 | | | | | | |
| fREML : -523.604, Scale est: 0.0246, N: 1262 | | | | | | |
| 10-fold cross-validation RMSE: 0.158 ± 0.01 | | | | | | |

**Table S16:** Summary of generalized additive model relating valley oak fitness to summer maximum temperature transfer difference, used for spatial prediction of future relative fitness. Model is a simplified version of Eq. 1b (Table S5) fitted to average relative fitness of localities instead of families. Values are also averaged across the two common gardens. The 2014 relative fitness term (which represents average height of all maternal trees at a locality) is used to represent the baseline relative fitness distribution in the spatial prediction models. Refer to Appendix S7, S9, and Methods, “Simulating the effects of assisted gene flow on future fitness” for more information.

| Component | Term | Est. | Std Error | t | p |  |
| --- | --- | --- | --- | --- | --- | --- |
| A. parametric coefficients | (Intercept) | -1.048 | 0.017 | -61.842 | < 0.001 | *** |
| Component | **Term** | **edf** | **Ref. df** | **F-value** | **p** |  |
| B. smooth terms | (Tdiff, average) | 2.491 | 3.105 | 11.412 | < 0.001 | *** |
|  | Smooth term (2014 relative fitness) | 1.505 | 1.864 | 4.052 | 0.0318 | * |
| Signif. codes: 0 <= '***' < 0.001 < '**' < 0.01 < '*' < 0.05 | | | | | | |
| Adjusted R-squared: 0.350, Deviance explained 0.380 | | | | | | |
| fREML : -130.895, Scale est: 0.027, N: 95 | | | | | | |
| 10-fold cross-validation RMSE: 1.41 ± 0.05 | | | | | | |

**Table S17**: Summary of geographic and climatic effects on cumulative relative fitness in 2024 at the Placerville garden. Predictor variables are scaled as Z-scores (“scale” function in base R) because of differences in their numeric ranges and scales.

|  | **Est.** | **Standard Error** | **t** | **p** |  |
| --- | --- | --- | --- | --- | --- |
| (Intercept) | 0.305 | 0.006 | 50.019 | < 0.001 | *** |
| scale(PC1) | 0.045 | 0.007 | 6.260 | < 0.001 | *** |
| scale(PC2) | -0.027 | 0.009 | -2.955 | 0.0032 | ** |
| scale(Latitude) | -0.004 | 0.016 | -0.285 | 0.7754 |  |
| scale(Longitude) | -0.024 | 0.015 | -1.586 | 0.1131 |  |
| *Signif. codes: 0 <= '***' < 0.001 < '**' < 0.01 < '*' < 0.05* | | | | | |
| Residual standard error: 0.1556 on 648 degrees of freedom | | | | | |
| Multiple R-squared: 0.08227, Adjusted R-squared: 0.0766 | | | | | |
| F-statistic: 14.52 on 648 and 4 DF, p: < 0.0001 | | | | | |

**Table S18**: Summary of geographic and climatic effects on cumulative relative fitness in 2024 at the Chico garden. Predictor variables are scaled as Z-scores because of differences in their numeric ranges and scales.

|  | **Est.** | **Standard Error** | **t** | **p** |  |
| --- | --- | --- | --- | --- | --- |
| (Intercept) | 0.403 | 0.006 | 69.128 | < 0.001 | *** |
| scale(PC1) | 0.020 | 0.007 | 2.911 | 0.0037 | ** |
| scale(PC2) | -0.025 | 0.009 | -2.851 | 0.0045 | ** |
| scale(Latitude) | -0.002 | 0.015 | -0.119 | 0.9054 |  |
| scale(Longitude) | -0.017 | 0.014 | -1.220 | 0.2230 |  |
| *Signif. codes: 0 <= '***' < 0.001 < '**' < 0.01 < '*' < 0.05* | | | | | |
| Residual standard error: 0.1467 on 630 degrees of freedom | | | | | |
| Multiple R-squared: 0.04182, Adjusted R-squared: 0.03573 | | | | | |
| F-statistic: 6.873 on 630 and 4 DF, p: < 0.0001 | | | | | |

#### **References for supporting information**

1. AdaptWest, P. (2022). Gridded current and projected climate data for North America at 1km resolution, generated using the ClimateNA v7.30 software (T. Wang et al., 2022). Data Basin adaptwest.databasin.org.

2. Bates, D., Mächler, M., Bolker, B. & Walker, S. (2015). Fitting linear mixed-effects models using lme4. *Journal of Statistical Software*, 67, 1–48.

3. Browne, L., Wright, J.W., Fitz-Gibbon, S., Gugger, P.F. & Sork, V.L. (2019). Adaptational lag to temperature in valley oak (*Quercus lobata*) can be mitigated by genome-informed assisted gene flow. *Proceedings of the National Academy of Sciences of the United States of America*, 116, 25179–25185.

4. Flint, L.E., Flint, A.L., Thorne, J.H. & Boynton, R. (2013). Fine-scale hydrologic modeling for regional landscape applications: the California Basin Characterization Model development and performance. *Ecological Processes*, 2, 25.

5. Gerst, K.L., Rossington, N.L. & Mazer, S.J. (2017). Phenological responsiveness to climate differs among four species of *Quercus* in North America. *Journal of Ecology*, 105, 1610–1622.

6. Girard, Q., Ducousso, A., de Gramont, C.B., Louvet, J.M., Reynet, P., Musch, B. *et al.* (2022). Provenance variation and seed sourcing for sessile oak (*Quercus petraea* (Matt.) Liebl.) in France. *Annals of Forest Science*, 79, 27.

7. Hijmans, R.J., Bivand, R., Pebesma, E. & Sumner, M.D. (2022). terra: Spatial Data Analysis.

8. Intergovernmental Panel on Climate, C. (2023). *Climate Change 2021 – The Physical Science Basis: Working Group I Contribution to the Sixth Assessment Report of the Intergovernmental Panel on Climate Change*. 1 edn. Cambridge University Press.

9. Karger, D.N., Conrad, O., Bohner, J., Kawohl, T., Kreft, H., Soria-Auza, R.W. *et al.* (2017). Climatologies at high resolution for the earth's land surface areas. *Sci Data*, 4, 170122.

10. Karger, D.N., Nobis, M.P., Normand, S., Graham, C.H. & Zimmermann, N.E. (2023). CHELSA-TraCE21k – high-resolution (1-km) downscaled transient temperature and precipitation data since the Last Glacial Maximum. *Climate of the Past*, 19, 439–456.

11. Mahony, C.R., Wang, T., Hamann, A. & Cannon, A.J. (2022). A global climate model ensemble for downscaled monthly climate normals over North America. *International Journal of Climatology*, 42, 5871–5891.

12. Pedersen, E.J., Miller, D.L., Simpson, G.L. & Ross, N. (2019). Hierarchical generalized additive models in ecology: an introduction with mgcv. *PeerJ*, 7, e6876.

13. Posit team (2025). RStudio: Integrated Development Environment for R. Posit Software, PBC Boston, MA.

14. R Core Team (2025). R: A Language and Environment for Statistical Computing. R Foundation for Statistical Computing Vienna, Austria.

15. Ramírez-Valiente, J.A., del Blanco, L.S., Alía, R., Robledo-Arnuncio, J.J. & Climent, J. (2022). Adaptation of Mediterranean forest species to climate: Lessons from common garden experiments. *Journal of Ecology*, 110, 1022–1042.

16. Rehfeldt, G.E., Ying, C.C., Spittlehouse, D.L. & Hamilton, D.A. (1999). Genetic responses to climate in *Pinus contorta*: niche breadth, climate change, and reforestation. *Ecological Monographs*, 69, 375–407.

17. Sáenz-Romero, C., Lamy, J.-B., Ducousso, A., Musch, B., Ehrenmann, F., Delzon, S. *et al.* (2017). Adaptive and plastic responses of Quercus petraea populations to climate across Europe. *Global Change Biology*, 23, 2831–2847.

18. St. Clair, J.B., Howe, G.T. & Kling, J.G. (2020). The 1912 Douglas-Fir Heredity Study: Long-Term Effects of Climatic Transfer Distance on Growth and Survival. *Journal of Forestry*, 118, 1–13.

19. Wadgymar, S.M., Sheth, S., Josephs, E., DeMarche, M. & Anderson, J. (2024). Defining fitness in evolutionary ecology. *Int J Plant Sci*, 185, 218–227.

20. Wang, T., Hamann, A., Spittlehouse, D. & Carroll, C. (2016). Locally downscaled and spatially customizable climate data for historical and future periods for North America. *PLoS One*, 11, e0156720.

21. Wickham, H. (2023). *modelr: Modelling Functions that Work with the Pipe*.

22. Wood, S.N. (2011). Fast stable restricted maximum likelihood and marginal likelihood estimation of semiparametric generalized linear models. *Journal of the Royal Statistical Society Series B: Statistical Methodology*, 73, 3–36.

23. Wood, S.N. (2017). *Generalized Additive Models: An Introduction with R, Second Edition*. 2 edn. Chapman and Hall/CRC, New York.

24. Wright, J.W., Ivey, C.T., Canning, C. & Sork, V.L. (2021). Timing of bud burst is associated with the climate of maternal origin in *Quercus lobata* progeny grown in a common garden. *Madroño*, 68, 443–449.
